# Multimodal spatiotemporal phenotyping of human organoid development

**DOI:** 10.1101/2022.03.16.484396

**Authors:** Philipp Wahle, Giovanna Brancati, Christoph Harmel, Zhisong He, Gabriele Gut, Aline Santos, Qianhui Yu, Pascal Noser, Jonas Simon Fleck, Bruno Gjeta, Dinko Pavlinić, Simone Picelli, Maximilian Hess, Gregor Schmidt, Tom Lummen, Yanyan Hou, Patricia Galliker, Magdalena Renner, Lucas Pelkmans, Barbara Treutlein, J. Gray Camp

**Affiliations:** Department of Biosystems Science and Engineering, ETH Zürich, Basel, Switzerland; Institute of Molecular and Clinical Ophthalmology Basel, Switzerland; Department of Ophthalmology, University of Basel, Basel, Switzerland; Department of Molecular Life Sciences, University of Zurich, 8057 Zurich, Switzerland; Roche Institute for Translational Bioengineering (ITB), Roche Pharma Research and Early Development, Roche Innovation Center Basel, Switzerland

## Abstract

Organoids generated from human pluripotent stem cells (PSCs) provide experimental systems to study development and disease. However, we lack quantitative spatiotemporal descriptions of organoid development that incorporate measurements across different molecular modalities. Here we focus on the retina and use a single-cell multimodal approach to reconstruct human retinal organoid development. We establish an experimental and computational pipeline to generate multiplexed spatial protein maps over a retinal organoid time course and primary adult human retina, registering protein expression features at the population, cellular, and subcellular levels. We develop an analytical toolkit to segment nuclei, identify local and global tissue units, infer morphology trajectories, and analyze cell neighborhoods from multiplexed imaging data. We use this toolkit to visualize progenitor and neuron location, the spatial arrangements of extracellular and subcellular components, and global patterning in each organoid and primary tissue. In addition, we generate a single-cell transcriptome and chromatin accessibility time course dataset and infer a gene regulatory network underlying organoid development. We then integrate genomic data with spatially segmented nuclei into a multi-modal atlas enabling virtual exploration of retinal organoid development. We visualize molecular, cellular, and regulatory dynamics during organoid lamination, and identify regulons associated with neuronal differentiation and maintenance. We use the integrated atlas to explore retinal ganglion cell (RGC) spatial neighborhoods, highlighting pathways involved in RGC cell death. Finally, we show that mosaic CRISPR/Cas genetic perturbations in retinal organoids provide insight into cell fate regulation. Altogether, our work is a major advance toward a virtual human retinal organoid, and provides new directions for how to approach disorders of the visual system. More broadly, our approaches can be adapted to many organoid systems.

## Introduction

Technologies to measure multiple molecular modalities in single cells are transforming our ability to explore developmental biology (Schier, 2020; Zhu et al., 2020). Transcriptomes and accessible chromatin can be profiled in thousands of cells per experiment (Schier, 2020; Zhu et al., 2020), and multiplexed imaging methods provide high-information content spatial registrations of tissues (Hickey et al., 2021). Within developing systems, single-cell sequencing and image-based measurements can be used to reconstruct cell state trajectories, which promise new insight into the differentiation dynamics across lineages and spatial domains. Applied to human stem cell-derived organoids (Chiaradia & Lancaster, 2020; Takebe & Wells, 2019), these technologies could be used to understand how molecularly-defined cell states relate to tissue structure and morphological development, and ultimately to create predictive virtual models of human disease. Indeed, it is a major goal in systems biology to generate in silico tissue models that incorporate increasing complexity, capturing multiple high-resolution length scales and include multiple cellular feature modalities (Camp et al., 2019; Rood et al., 2019). A major challenge to achieve this goal is to integrate multimodal measurements across multiple length scales in meaningful ways to reveal the mechanisms by which scale-crossing effects drive tissue development and morphogenesis. In particular for human development, where embryonic samples are scarce, there is the additional challenge to achieve this with organoid models, which often lack stereotypic organization, with substantial heterogeneity within and between organoids. There is thus a large unmet need to implement and integrate multimodal technologies in developmentally dynamic and human-relevant model systems (Hao et al., 2021).

Retinal organoids generated from induced pluripotent stem cells (iPSCs) offer an inroad into studying human retina development, identifying mechanisms of disease, and facilitating the discovery of new treatments (Brancati et al., 2020). From the breakthrough discovery that optic cups spontaneously self-organize in three-dimensional PSC-derived cultures in vitro (Eiraku et al., 2011; Nakano et al., 2012; Sasai, 2013), multiple different protocols to generate human retinal organoids have been developed (Afanasyeva et al., 2021; Cowan et al., 2020; Zhong et al., 2014). Retinal organoids are composed of diverse neural cell types - rod and cone photoreceptors (PRs), bipolar cells (BCs), horizontal cells (HCs), amacrine cells (ACs), retinal ganglion cells (RGCs), and Müller glia (MG). These cells self-organize into a stereotypical laminar structure with outer and inner plexiform layers and outer (PR nuclei) and inner (AC, BCs and HC) nuclear layers (ONL and INL), together with RGCs in the retinal ganglion cell layer. Remarkably, immunohistochemical analysis and single-cell transcriptome characterization of developed organoids has confirmed their high similarity to the primary human retina (Cowan et al., 2020; Lu et al., 2020; Sridhar et al., 2020), providing a foundation for understanding human retina development and disease in vitro. Retinal organoids develop over the course of months, reaching a maximal in vitro maturation state around the age of 30 weeks with current methods (Cowan et al., 2020). Multiple studies have shown the potential value of retinal organoids for modeling human vision disorders (Dulla et al., 2018; Khan et al., 2020; Kruczek et al., 2021; Parfitt et al., 2016). However, differentiation trajectories and morphological heterogeneity has yet to be quantitatively evaluated and we lack a reconstruction of the gene regulatory networks that underlie differentiation of each neuronal and glia cell type within the organoid tissues. More generally, we lack a foundation to integrate high-resolution and high-information content imaging data of tissue organization with sequencing data to assess phenotypes within these complex organoid tissues.

Here, we established an experimental pipeline for performing iterative indirect immunofluorescence imaging (4i) (Gut et al., 2018) on histological sections of retinal organoids at high spatial resolution, and develop a computational approach for inferring spatial developmental dynamics from multiplexed protein maps of heterogeneous organoids. We combine this with a dense scRNA-seq and scATAC-seq dataset covering 6 to 46 weeks of development which can reconstruct differentiation trajectories into each of the major neuronal and glial lineages. Integration of all data modalities provides a first-of-its-kind digital representation of human retinal organoid development. The digital organoid can be used to explore spatial interactions over time, and predict gene regulatory modules underlying retinal neurogenesis. We perform follow-up in organoid mosaic perturbations of selected transcription factors (TFs) and focus on the Orthodenticle Homeobox 2 (OTX2) regulon required for human retinal neurogenesis (Ghinia Tegla et al., 2020; Nishida et al., 2003; Wang et al., 2014). Altogether, our work is a major advance toward a virtual human retinal organoid, and our approaches can be adapted to other developing organoid or other model systems.

## Results

### A 4i atlas of human retinal organoids and adult retina tissue

To establish a spatial retinal organoid reference map, we applied 4i covering a time course of human retinal organoid development (6, 12, 18, 24, 39 weeks; 2-4 organoids per time point, Table S1) as well as an adult primary human retina (Fig. 1a). For each retinal organoid and the adult tissue, we performed multiplexed immunohistochemistry on 3μm thick formalin fixed paraffin embedded (FFPE) sections (1-4 sections per sample, Table S1)(Fig. 1b). We generated tiled images at 40x magnification with a high numerical aperture silicone oil objective to cover length scales from the millimeter to the nanometer scale (pixel size 0.1625 μm) (Fig. 1c). We established a panel of 63 antibodies (ABs) covering major retinal cell types, subcellular compartments, morphological structures and signaling pathways split into three color channels (Table S1). We obtained strong and reproducible signals over 21 imaging cycles, with usable data from 52 antibodies and a nuclei stain (Extended Data Fig. 1a,b). To achieve multiplexing, we aligned images (Marstal et al., 2016) across all cycles, allowing simultaneous display of any protein stain (Extended Data Fig. 1c-f, 2a,b). In our experimental design, we interspersed unstained cycles to subtract cycle-specific backgrounds, permuted ABs to assess influence of AB order, and performed elution controls (Methods). These technical assessments confirmed that the staining patterns passing quality control were robust (Extended Data Fig. 2c-f). Altogether, this resulted in an expression matrix with over 900 million pixels registering 53 fluorescence intensity measurements from a retinal organoid developmental time course comprising 42 retinal organoid sections and an adult human retina sample.

**Fig. 1.**
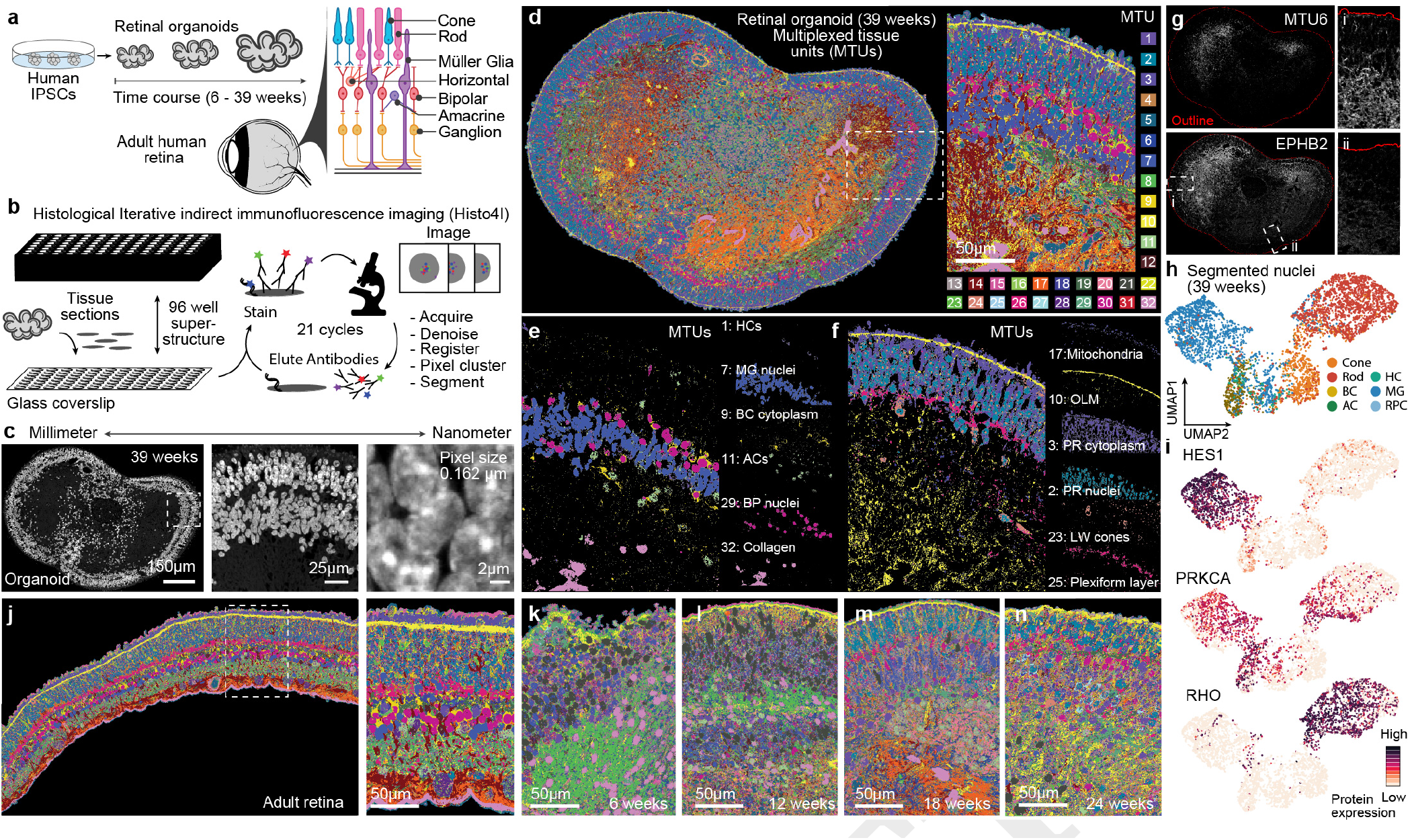
Highly multiplexed immunohistochemistry reveals scale-crossing features of developing retinal organoids and primary human retina. a) We performed 4i on tissues of a time course of retinal organoid development (6, 12, 18, 24, 39 weeks) as well as adult retina tissue. Schematic shows retinal organization and broad cell types. b) Schematic of the 4i methodology. FFPE tissue sections were placed on a 96-well format coverslip that was subsequently attached to a 96-well superstructure allowing immunohistological treatment followed by imaging and AB elution. Here we performed 21 immunohistochemical staining cycles. Images were acquired at 40X magnification with a high numerical aperture silicone oil objective, tiled across the tissue. c) Images showing overview of a Hoechst stain of a 39 week organoid section and progressive magnifications from millimeter to nanometer scales. d-f) Example of pixel clustering of a 39 week organoid section. d) 32 global MTUs resolve the tissue structure with image quality and label biological structures in individual samples. e-f) Biological structures identified by unique MTUs include horizontal cells (HCs) bipolar cell (BC) cytoplasm and nuclei, amacrine cells (ACs) and structural elements such as collagen rich areas (e) and peripherally located structures such as Mitochondria, the outer limiting membrane (OLM) photoreceptor (PR) cytoplasm and nuclei, long wave (LW) cones and the plexiform layers (f). g) MTU 6 (top) is enriched for EPHB2 protein expression (bottom) and non uniformly distributed in the organoid section (outlined by red line). Insets show two regions with high (i) and low (ii) detection of EPHB2 fluorescence immuno histochemical signal. h) Heterogeneity analysis of nuclei with UMAP projection based on protein features and colored by labels transferred from sequenced cells. All major types are identified including retinal ganglion cells (RGCs), horizontal cells (HCs), cones, amacrine cells (ACs), rods, bipolar cells (BPs), müller glia (MG). i) Feature plots highlighting median signal level per nucleus of HES1, identifying nuclei located in the INL, PRKCA, enriched in Rod BP cells and RHO, identifying cells located in the ONL.

For all tissues and time points, we established a multi-scale analysis pipeline that includes an unsupervised machine learning-based clustering of pixels from their 53-plex intensity profile (termed multiplexed tissue units, MTUs) (Gut et al., 2018), nuclei segmentation, assessment of nuclei heterogeneity, spatial arrangement from protein intensities and MTU distributions (Methods). MTUs can be generated for single samples or for the entire time course (global MTUs, Methods) (global MTUs, Methods) (Extended Data Fig. 3a-f). Throughout the manuscript we analyze global MTUs. Focusing on an exemplary 39 week organoid, this revealed 32 MTUs that provide a detailed characterization of the spatial organization of the tissue (Fig. 1d). Hierarchical clustering and heatmap visualization of average protein expression within each MTU highlights how each protein stain associates with each MTU (Extended Data Fig. 4a). We find that the MTU patterns are similar across different sections from the same organoid, as well as between organoids of the same time point (Extended Data Fig. 4b). This indicates that MTUs provide a meaningful approach to quantify principles of tissue organization and composition in an unbiased manner, which is robust to the morphological heterogeneity observed within and between organoids. We observed that certain MTUs distinguish different cell types, whereas others resolve subcellular and tissue structures (Fig. 1e,f), illustrating their versatile multi-scale nature. For example, MTU 9 and 29 segment a subset of bipolar cell nuclei and cytoplasm, respectively, MTU 17 resolves mitochondria, MTU 10 highlights the outer limiting membrane, and MTU 25 shows the outer and inner plexiform layers. MTU 6 identified a surprising feature marked by Ephrin type-B receptor 2 (EPHB2) (Fig. 1g). EPHB2 functions in axon guidance and is asymmetrically expressed in retinal neurons demarcating dorsal and ventral domains of the developing mammalian retina (Birgbauer et al., 2000), suggesting that dorsoventral patterning domains can emerge during human retinal organoid development. Nuclei variance analysis in the protein expression feature space and visualization in a Uniform Manifold Approximation and Projection (UMAP) embedding, revealed 8 molecularly distinct nuclei clusters (Extended Data Fig. 4c,d), and comparison of protein expression with single-cell transcriptomes (Cowan et al., 2020) enabled cell type assignment to each nuclei cluster (Fig. 1 h,i; Extended Data Fig. 4e,f). The major retina cell types could be identified including photoreceptors, RGCs, HCs, ACs, BCs, and MG. Over the time course and in the adult primary tissue, we also observed striking progression of MTU and nuclei organization patterns further showcasing the robustness of the methods and the scale-crossing richness of the dataset for exploring developmental phenomena (Fig. 1j-n, Extended Data Fig. 3). We provide a web application (EyeSee4is, https://eyesee4is.ethz.ch/) to facilitate access to the imaging data and computed features over the time course. Altogether, these data establish 4i on tissues as a flexible, robust, sensitive, and high-dimensional method to describe organoid cell composition and structure based on protein measurements.

### Laminar structure trajectory analysis illuminates spatiotemporal dynamics of retinal layer formation

To study how the laminar structure in retinal tissue emerges, we developed a computational approach to reconstruct organoid laminar structure dynamics from multiplexed imaging data. This method (Laminator) establishes a contour around the organoid, segments and vertically orients adjacent laminar windows circumference-spanning each organoid, quantifies signals across laminar windows, analyzes laminar window heterogeneity, and applies graph embedding and diffusion analysis for trajectory reconstruction (Fig. 2a). We measured immunofluorescence intensities, MTU distributions, and nuclei features in an inner-to-outer axis per laminar window, clustered the features per oriented laminar window, and visualized relationships within and across organoids using a UMAP embedding (Extended Data Fig. 5a-c). From this analysis we could distinguish various structural components of the tissue, such as highly organized nuclear layers, disorganized zones, non-retinal regions, and aggregates of retina pigmented epithelium (RPE). We selected retinal regions and used a force-directed graph to analyze the relationships between clusters across the time points (Fig. 2b), and applied diffusion analysis to establish a trajectory of retinal neuron layer development (Fig. 2c; Extended Data Fig. 5d, e). The reconstructed pseudotemporal trajectory reflected a temporal progression (Fig. 2d), and enabled us to observe several interesting aspects of human retinal organoid development preceding retinal lamination and spanning photoreceptor maturation (Fig. 2e; Extended Data Fig. 5f-h). In early phases of the trajectory, there were abundant cells in the G2/M phase of the cell cycle marked by MKI67 and PCNA, and associated nuclei localized to outer surfaces and exhibited elongated shapes (Fig. 2e; Extended Data Fig. 6a). Progenitor cells begin to differentiate into neurons and subsequently the plexiform layers emerge, becoming stratified between nuclei layers (Fig. 2e,f). In a later stage, photoreceptors develop and differentiation of neuronal and glial cells is established (Extended Data Fig. 5h, 6b,c). Overall, there is a clear and consistent pattern across diverse signals that retinal organoids increase in similarity to the primary retina organization over the time course (Fig. 2g), consistent with scRNA-seq data (Cowan et al., 2020). We note that this reconstruction can be used to analyze the dynamic location of subcellular structures such as mitochondria and P-bodies, and tissue structures such as collagen and axonal fibers (Extended Data Fig. 5h, 6b,c). Altogether, these data provide a high-information content spatial representation from protein measurements of human retinal organoid laminar developmental dynamics.

**Fig. 2.**
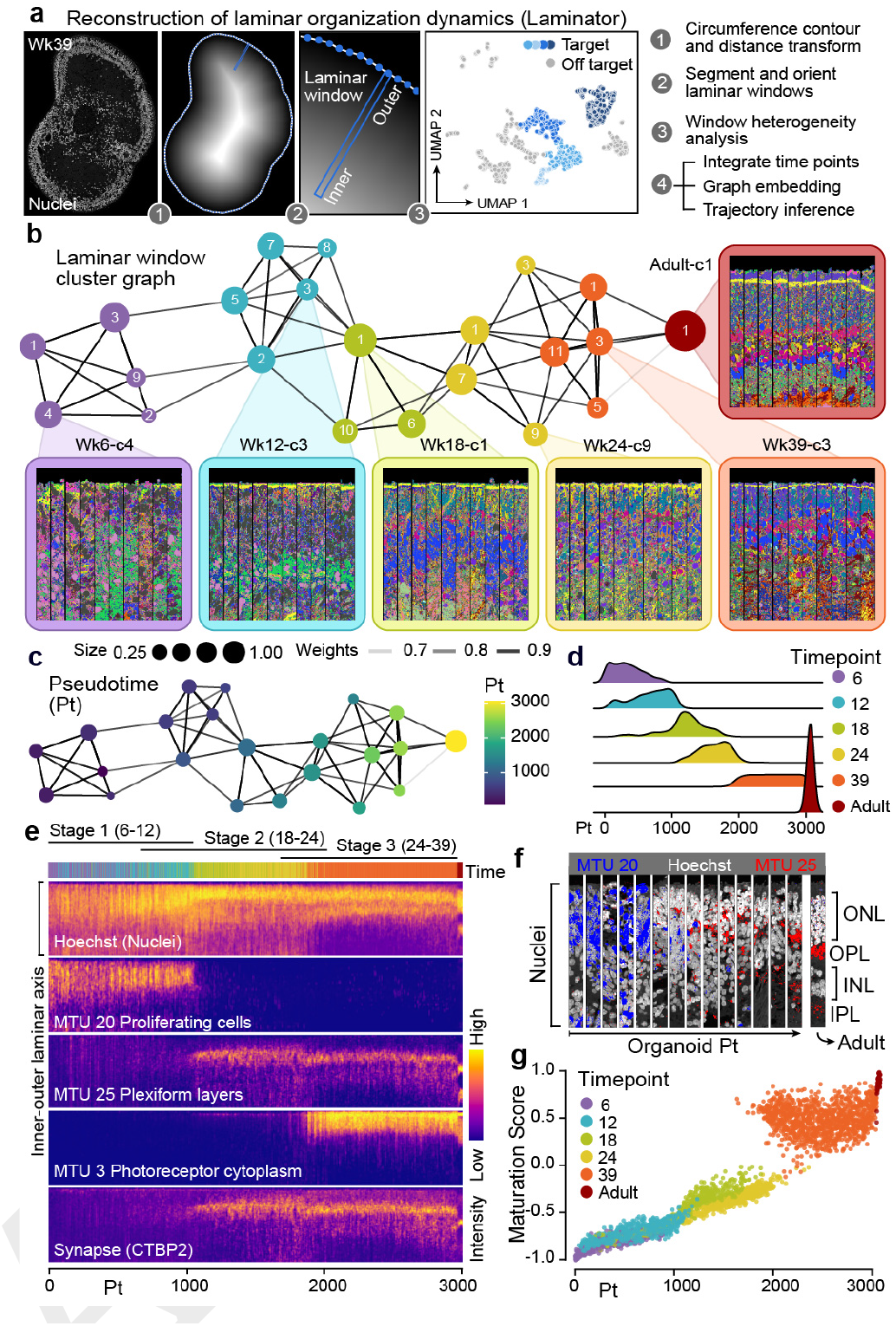
Analysis of laminar structure enables trajectory reconstruction and illuminates spatiotemporal dynamics of retinal organoid formation. a) Schematic overview of the Laminator algorithm developed for laminar window segmentation, vertical orientation, and trajectory inference. b) Force-directed graph embedding of laminar window clusters (numbered) colored by time point. Node size represents the fraction of laminar windows within a time point. Insets colored by timepoints show representative oriented laminar windows per cluster. c) Graph with laminar window clusters colored based on pseudotime from diffusion analysis. d) Density plot showing laminar window proportion along the pseudotime, grouped by time point. e) Heatmap showing fluorescence intensity measurements (Hoechst and CTBP2) or MTU intensity profiles (MTU 20, 25, 3) along the inner-outer laminar axis across oriented laminar windows ordered by pseudotime. f) Representative oriented laminar windows from multiple positions along the pseudotime course showing nuclei location (Hoechst, white), proliferating cells (MTU 20, blue), and plexiform layers (MTU 25, red) along the inner-outer laminar axis. g) Scatter plot showing signal similarity of each laminar window to adult laminar windows over pseudotime. Dots are colored by time point.

### Time course single-cell transcriptome and accessible chromatin profiling identifies gene regulatory networks underlying human retinal organoid development

In order to provide multi-omic resolution to human retinal organoid developmental dynamics, we performed scRNA-seq and scATAC-seq from the same cell suspension across a time course (6-46 weeks) of human retinal organoid development (Fig. 3a). The dataset incorporates 22 time points from 4 human induced pluripotent stem cell (iPSC) lines, including iPSC lines with stable integration of doxycycline-inducible Cas9 in the AAVS1 safe harbor locus (iCas9), and previously published scRNA-seq datasets (Cowan et al., 2020) (Table S2). We constructed ‘multi-omic metacells’ containing information on both transcriptome and chromatin accessibility using minimum-cost, maximum-flow (MCMF) bipartite matching (Stark et al., 2020) within canonical correlation analysis (CCA) space (Stuart et al., 2019) (Extended Data Fig. 7a). The metacells were integrated using cluster similarity spectrum (CSS) (He et al., 2020) and the integrated data was visualized using a UMAP embedding. In addition, we performed Multiome measurements from the same cell (10x Genomics) at key developmental time points (15 and 36 weeks), incorporated the cells into the integration, and used the Multiome data to assess the overall integration (Extended Data Fig. 7b-h). Altogether, the integration revealed a continuous distribution of cell states through the time course, with rods, cones, BCs, ACs, HCs, RGCs, RPE, and MG annotated (Fig. 3b). The high-dimensionality of the data could be used to identify marker genes and gene regulatory regions for the different cell types (Fig. 3c,d, Table S3). Indeed, the cell type-specific gene regulatory regions overlap with cell type-specific promoters and enhancers (Extended Data Fig. 7i-o), which may be useful to design cell type-specific drivers for gene therapy (Jüttner et al., 2019). These data provide a high-resolution feature assessment of human retinal organoid development from multipotent progenitor states.

**Fig. 3.**
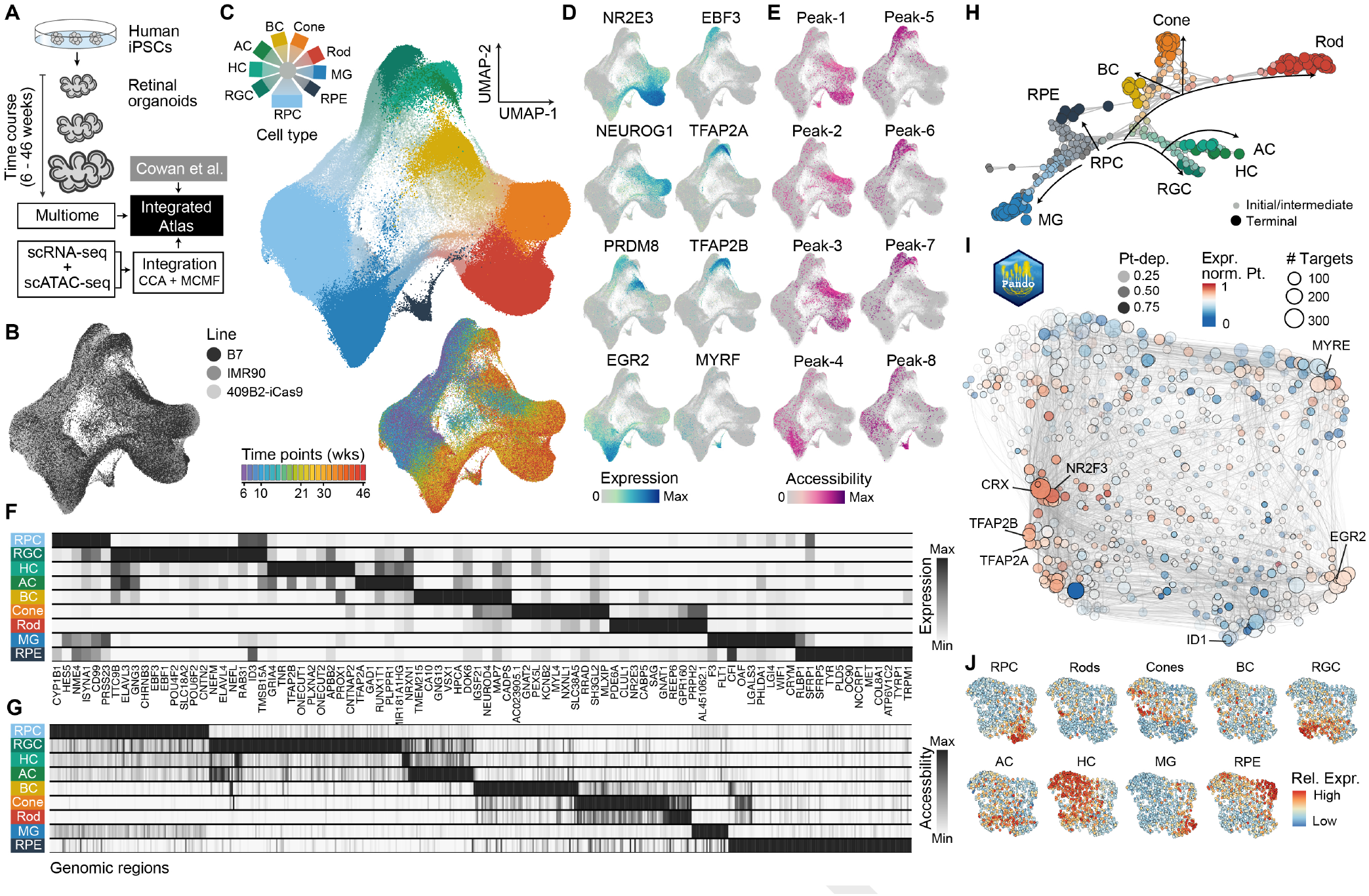
Time course single-cell multiomic data identifies gene regulatory networks underlying human retinal organoid development. a) Paired scRNA-seq and scATAC-seq data were performed on a time course of retinal organoid development. Multiome data was also acquired and used to assist with data integration. Together with previously published scRNA-seq data 14, scRNA-seq and scATAC-seq data were combined into metacell representations containing both modalities using canonical correlation analysis (CCA) and minimum cost maximum flow (MCMF). b) UMAP embedding of metacells colored by annotated cell type (left), time point (top right), or iPSC line (bottom right). c) Feature plots showing cell type marker gene expression (left, blue scale) or chromatin accessibility (right, pink scale). d) Heatmaps showing average expression of representative marker gene expression (top) or chromatin accessibility (bottom) for each major cell type. e) Branch visualization in a force-directed layout, with circles representing high-resolution clusters with both RNA and access features colored by assignment. f) UMAP embedding of the inferred TF network based on co-expression and inferred interaction strength between TFs. Color and size represent expression weighted pseudotime of TF regulator and pagerank centrality of each module. g) TF network colored by expression enrichment for different cell types.

We next reconstructed the cell and gene regulatory networks (GRNs) that underlie human retinal development. We used RNA velocity (La Manno et al., 2018) and CellRank (Lange et al., 2022) to generate a terminal fate transition probability matrix based on transcriptomes, which we used to construct a differentiation graph of high-resolution metacell clusters and assign branch identities. The graph, presented by a force-directed layout, reveals diversification of retinal cell types over the organoid time course (Fig. 3e). We used Pando (Fleck et al., 2021), to infer sets of positively or negatively regulated target genes (gene modules) as well as regulatory genomic regions (regulatory modules) for each annotated TF (Fig. 3f) and visualized module expression in a UMAP embedding. Feature plots reveal groups of TFs that associate with the development of specific cell types (Fig. 3g), including RGC development (POU4F2, ATOH7), HC/AC/BC differentiation (TFAP2A, TFAP2B, PRDM8), and photoreceptor diversification (rod NR2E3; cone NEUROG1, both: CRX). Globally, this GRN shows that regulatory region accessibility and TF expression track with stages of retinal organoid development and segregate during neuron diversification.

### Multimodal data integration provides a digital developmental map of retinal neurogenesis

We next sought to integrate the sequencing and multiplexed imaging data in order to generate a multimodal map of human retinal organoid development. We subsetted metacells from each sequencing time point, performed high-resolution clustering in the transcriptome space, and predicted 4i nuclei type based on correlation between transcript and protein expression (Fig. 4a). In this way, the transcriptome and accessible chromatin modalities could be integrated with spatially localized nuclei in the 4i dataset, such that each nucleus contains chromatin, mRNA, protein, and spatial features (multimodal nuclei, Fig. 4b, Extended Data Fig. 8a). Focusing on the developed organoid (39 weeks), we highlight CRX and HES1 expression in photoreceptors and MG respectively (Fig. 4c), as well as rod and MG-specific gene regulatory regions (Fig. 4d). Applied to all sections and time points, we could map protein abundance, nuclei location, transcriptome, and chromatin accessibility in all tissues, facilitating the spatial exploration of diverse data modalities across time (Fig. 4e, Extended Data Fig. 8). For example, we show that VSX2 is initially highly expressed in progenitor cells localized towards the outer portion of the early organoid (6-12 weeks), and it becomes restricted to bipolar cells localized in the INL in developed organoids (39 weeks) (Fig. 4f, g). We could resolve the temporal emergence of each neuronal class, observing the transition from multipotent progenitor cells that were distributed throughout the lamina into intermediates and differentiated types that began to stratify over time to the INL and ONL (Fig. 4h, i). This analysis highlighted the emergence and disappearance of RGCs (marked by POU4F1), a known deficiency of current retinal organoid protocols (Fig. 4h-j) (Capowski et al., 2019; Cowan et al., 2020; Zhong et al., 2014).

**Fig. 4.**
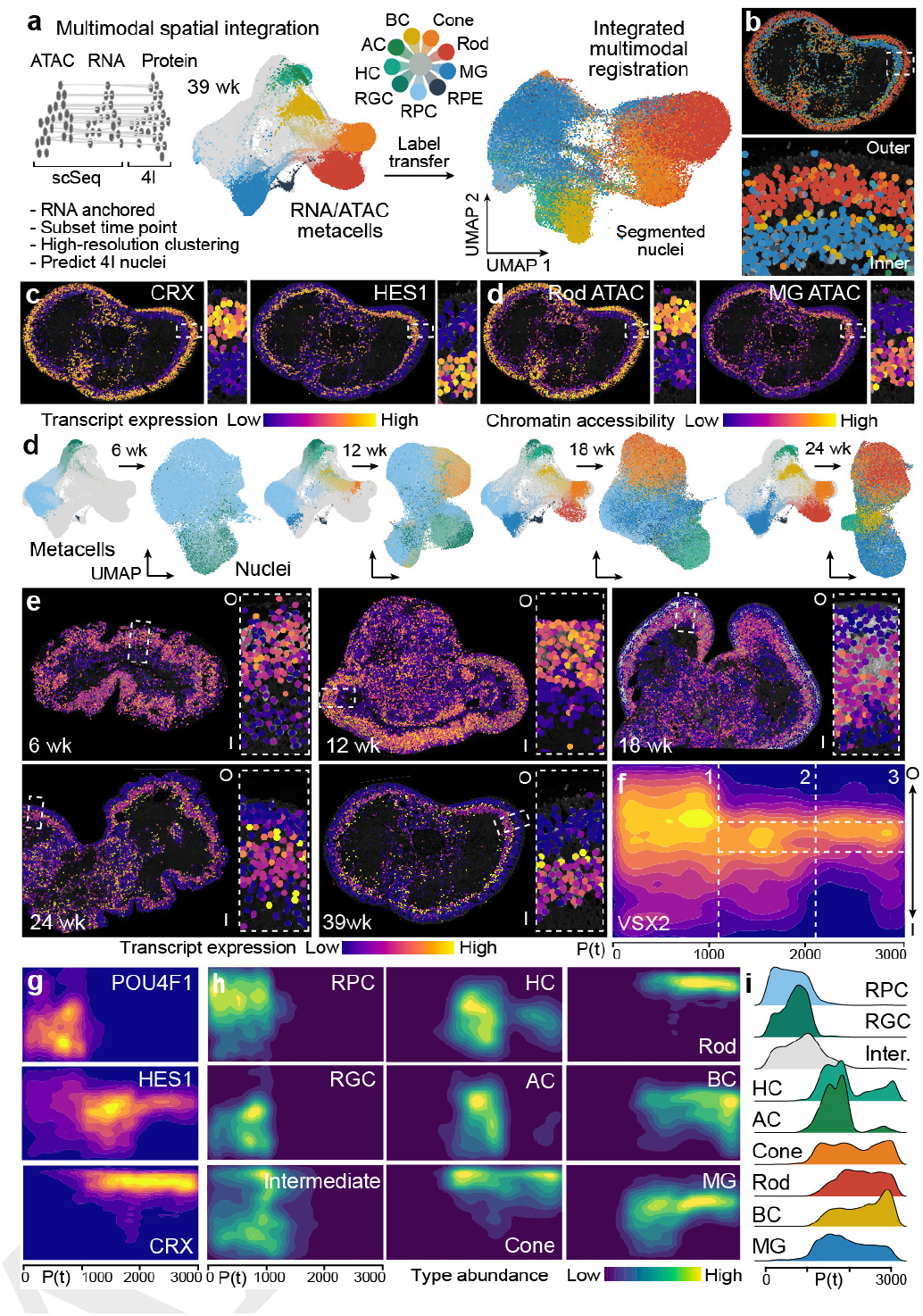
Multimodal data integration into a spatiotemporal digital organoid representation provides a virtual map of retinal neurogenesis. a) Schematic for integrating accessible chromatin, transcriptome, and protein modalities into spatially resolved and segmented nuclei. High-resolution clusters were generated from scSeq data (RNA/ATAC metacells) of the closest matching time points to the imaging data. Label transfer was predicted based on correlation of RNA and protein features in sequenced cells and imaged nuclei, respectively. Left UMAP shows time course metacell embedding based on transcriptome and colored by cell type with 39 week cells highlighted. Right UMAP shows nuclei embedding based on protein features colored according to label transfer from the transcriptome space. b) Overview and laminar zoom of a reference 39 week retinal organoid colored based on nuclei-type assignment from the label transfer. c-d) Reference 39 week organoid nuclei colored based on RNA expression (c) or chromatin accessibility (d) of markers for photoreceptors (CRX, c; chr17-81655189-81657223, d) or muller glia (HES1, c; chr11-126082119-126083088, d). e) Multimodal integration across the other time points. Left metacell embedding colored based on indicated time point. Right nuclei UMAP colored by label transfer from the transcriptome space. f) Time course retinal organoids colored based on VSX2 transcript expression. Boxed inset shows zoom with inner (i) to outer (o) orientation. g) Heatmap shows VSX2 expression densities along the inner-outer and pseudotime axes. Dotted lines demarcate stages (vertical) and INL (horizontal). h) Heatmap showing expression of retinal ganglion cells (RGC, POU4F1), müller glia (MG, HES1), and photoreceptor (CRX) markers along the inner-outer and pseudotime axes. i) Heatmap showing nuclei type abundance densities for the major annotated retinal organoid cell types/states along the inner-outer and pseudotime axes. j) Density plots showing proportion of each annotated nuclei type over the trajectory.

We used the extended dimensionality of the integration to identify transcriptome features that associate with spatial features. We first focused on larger spatial domains. Previous analyses identified MTU 6 in a 39 week organoid that was distinguished by EPHB2 protein expression (Fig. 1g), a delineator of retinal dorsal-ventral patterning. We observed that multimodal nuclei in the integration also showed spatial heterogeneity in EPHB2 transcript abundance (Extended Data Fig. 9a). We searched the multimodal nuclei for transcripts that spatially correlated and anticorrelated with EPHB2 expression. This analysis identified gene sets enriched in nuclei in EPHB2-high and EPHB2-low domains, and these genes had high expression in MGs and BCs and lower expression in photoreceptors (Extended Data Fig. 9b,c). We performed gene ontology (GO) term enrichment analysis of genes positively or negatively spatially correlated with EPHB2. This revealed general sensory or neuronal terms for genes negatively spatially correlated with EPHB2 while terms of genes positively spatially correlated with EPHB2 related to metabolism and development suggesting that cells high in EPHB2 might be developmentally active (Table S4, Extended Data Fig. 9d,e). These data support our previous observation that patterning domains can emerge in retinal organoids, which have an impact on longer term expression patterns that emerge after weeks in culture.

We next established a neighborhood analysis to explore local, microenvironmental variation during cell type differentiation. We segmented cell neighborhoods through a 6.5μm radial extension from each nuclei centroid, and then searched for heterogeneity among neighborhoods based on annotated features (e.g. MTUs, protein, RNA, chromatin access) (Extended Data Fig. 10a). We applied this pipeline to understand the loss of RGCs, as RGCs emerge at 6 weeks, become abundant at 12 weeks, but are nearly absent at 18 weeks of development. Clustering and visualizing RGC neighborhoods from 12 week organoids based on MTU profiles in a UMAP embedding identified significant heterogeneity among the RGC neighborhoods (Extended Data Fig. 10b,c). Inspection of RGC neighborhoods on the image, revealed differential location patterns between many of the clusters (Extended Data Fig. 10d). Interestingly, certain RGC neighborhoods were localized within the interior of the organoid, and nuclei within these neighborhood clusters exhibited cell death features, including intense Hoechst staining and nuclei fragmentation (Crowley et al., 2016), and other protein and MTU characteristics (Extended Data Fig. 10e-g). We explored heterogeneity in the transcriptome space, and identified two major RGC types, distinguished by the presence and absence of POU4F1 expression (Extended Data Fig. 10h). Sub-clustering revealed 11 clusters, and differential expression analysis between clusters highlighted cluster 6 as having a strong signature of apoptosis (Extended Data Fig. 10i-j). Gene ontology analysis revealed pathways, and specific genes, that may be involved in RGC preservation or induction of cell death, which have implications for human vision disorders such as glaucoma (Almasieh et al., 2012; Vrabec & Levin, 2007) (Table S5). Altogether, these data and analyses showcase how the digital retinal multimodal map can be used to explore gene regulation and spatial feature co-variation, and cell neighborhood analyses can be performed for any cell type or spatial domain.

### Mosaic perturbation of transcription factor regulomes provides insight into cell fate control in developing human retinal organoids

The GRN analyses and integrated multimodal map illuminated TFs that are central regulators of development. To begin to understand how TFs regulate retinal cell type identity in human tissues, we established a pooled loss of function (LOF) experiment based on the CROP-seq protocol (Datlinger et al., 2017; Fleck et al., 2021) in developed retinal organoids (Extended Data Fig. 11a). We targeted five TFs that are important for retinal development and expressed dynamically over the organoid developmental time course (OTX2, NRL, CRX, VSX2, and PAX6) (Swaroop et al., 2010), Extended Data Fig. 11b,c; Fig. 5a, 4f). We found OTX2, CRX and NRL are expressed in rods, cones, BCs and the RPE (Extended Data Fig. 11c). VSX2 and PAX6 are expressed in RPCs and their expression is maintained in BCs and RGCs/ACs/HCs respectively (Extended Data Fig. 11c). To perform the LOF experiment, we established an inducible Cas9 nuclease (iCas9) line, and validated that it produces each of the major retina neuronal cell types (Fig. 3b, d). We generated a lentiviral library containing a GFP reporter (Fleck et al., 2021) and targeting guides against OTX2, NRL, CRX, VSX2, and PAX6 and a non-targeting guide as control. The iCas9 line was used to generate retinal organoids, which were infected with the pooled lentiviral library at 19 weeks of development. At 3-5 weeks post-infection, GFP+ cells were sorted and used for scRNA-seq (10x Genomics) and target amplicon sequencing. Based on RNA expression measurements we generated a UMAP embedding, analyzed cell type heterogeneity, and annotated the major cell types recovered in the experiment (Extended Data Fig. 11d). We performed expression module analysis (Jin et al., 2020) and found that OTX2 LOF cells showed the strongest effect with OTX2 module genes being significantly misregulated (Extended Data Fig. 11e-g). We therefore focused subsequent detailed analyses on the effect of perturbation of the OTX2 regulon.

**Fig. 5.**
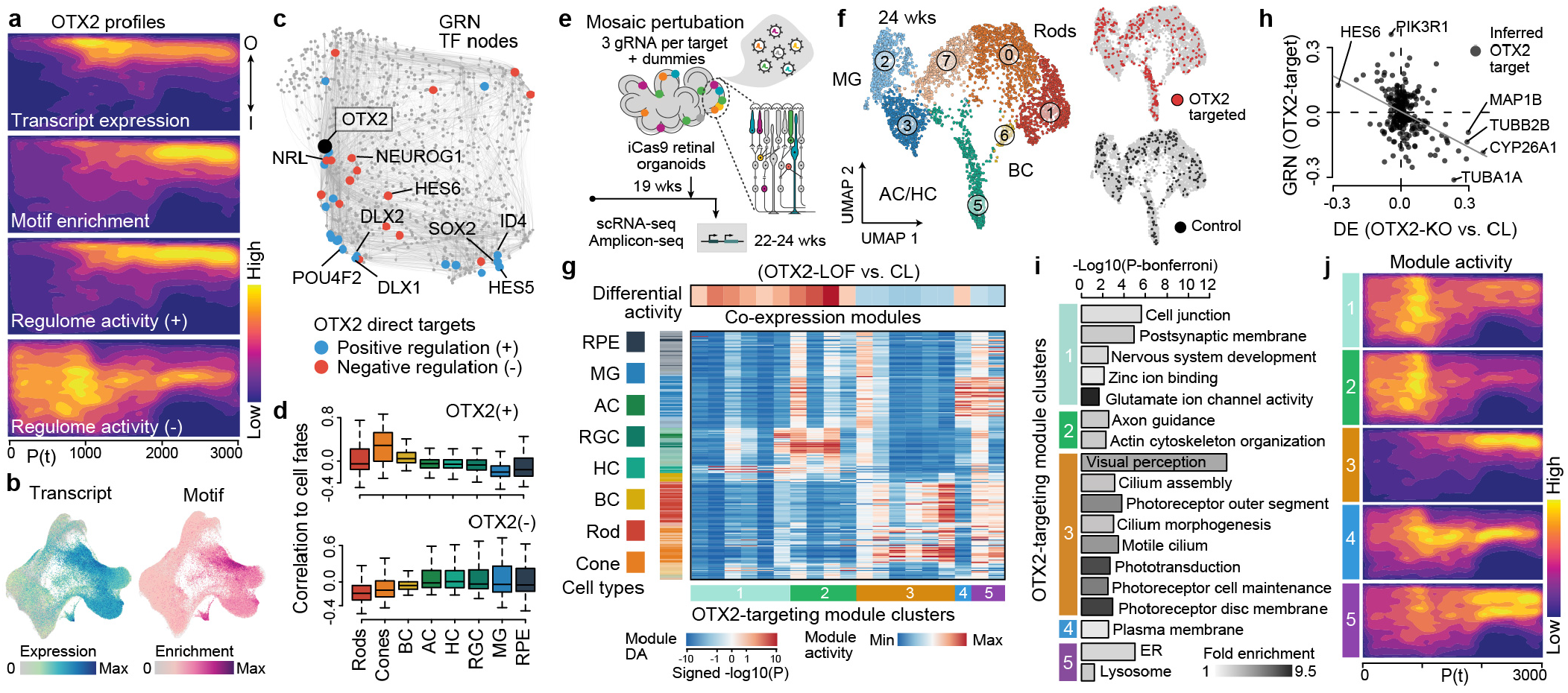
Perturbation of the OTX2 regulome has different effects across the developing retinal cell types. a) Heatmap shows OTX2 transcript expression, motif enrichment, positive regulome (+), and negative regulome (−) densities along the inner (I) to outer (O) and pseudotime axes from the reconstructed multimodal map. b) Transcriptome-based UMAP metacell plot showing OTX2 expression (left) or motif enrichment within accessible chromatin peaks (right). c) Global TF gene regulatory network (GRN) highlighting OTX2 (black node) and the predicted positively (blue) and negatively (red) regulated genes within the inferred OTX2 regulon. d) Boxplots show the distribution of OTX2 positively (top, +) or negatively (bottom, −) regulated targets within the GRN based on the expression correlation of each target to different retinal cell fates. e) Schematic of single-cell perturbation experiment using the CROP-seq method. Three gRNAs targeting OTX2 and 4 other TFs (see extended data) were used together with a random non-targeting gRNA (dummy). Retinal organoids were infected with gRNA-containing lentiviruses at 19 weeks, and scRNA-seq and amplicon-seq were performed on suspensions at 22-24 weeks. f) UMAP projection colored by annotated cell type (left) or by cells with OTX2 (top right, red) or dummy (bottom right, dark grey) gRNAs detected. Grey cells represent unknown or other targeted TFs. g) Heatmap showing gene expression modules (columns) and their activity in cell clusters (rows). Left sidebar shows cluster type. Bottom sidebar shows module clusters. Top side bar shows the differential module activity for the OTX2 gRNA cells relative to dummy control. h) Scatter plot showing the relationship between differential expression between OTX2 loss of function and control (x axis) and predicted directionality in the OTX2 GRN targets (y axis). i) Gene ontology enrichments for modules that are significantly affected by OTX2 loss of function. j) Heatmap shows the module activity scores across the retinal organoid spatiotemporal multimodal map.

We first explored OTX2 in our digital organoid map. The OTX2 regulon is distinguished by predicted positive regulation of genes enriched in rods, cones and BCs relative to the other retinal cell types, with the OTX2 motif being enriched within accessible chromatin of these cell types (Fig. 5a,b). Within the global GRN, we find that among direct and positively regulated OTX2 targets are NRL, NEUROG1, and HES6, which are TFs that are also expressed in photoreceptors or BCs. Conversely, direct negative targets are DLX1/2, HES5, and POU4F2, which are TFs enriched in fates that are predicted to be negatively regulated by OTX2 (Fig. 5c,d) (Ghinia Tegla et al., 2020). The CROP-seq experiment provided sufficient OTX2 gRNA detection across the different retinal cell types to assess the impact of predicted LOF on each cell type expression module (Fig. 5e,f). For OTX2-LOF, we found that RGC, HC, and AC expression modules were enriched, and photoreceptor and BC expression modules were depleted in comparison to control cells (Fig. 5g). In addition, there was a correlation between GRN OTX2 target predicted regulation directionality, and differentially expressed genes between OTX2-LOF and control cells (Fig. 5h). The most depleted expression module in the OTX2-LOF condition (module 3) had ontology enrichments for many aspects of visual perception, consistent with the role of OTX2 in maintaining photoreceptor identity and the critical role of OTX2 in the development and maintenance of the human visual system (Fig. 5i). Finally, we highlight the activity of these OTX2 regulated modules over the digital organoid map, showing that module 3 emerges temporally upon photoreceptor differentiation as they localize to outer positions within the lamina (Fig. 5j). These findings are consistent with previous results in non-human model systems showing that OTX2 governs sister fate choices in the developing retina, particularly directing photoreceptor and BC programs, while inhibiting the AC/HC/RGC programs (Ghinia Tegla et al., 2020; Nishida et al., 2003; Sato et al., 2007). In addition, we looked at how each targeted TF in the CROP-seq affects the regulome of the other TFs targeted in the CROP-seq (Extended Data Fig. 11h). Interestingly, this reveals that OTX2-LOF induces a strong effect on the PAX6 (Ghinia Tegla et al., 2020; Nishida et al., 2003) and CRX regulomes (Nishida et al., 2003). Since it is currently unknown how OTX2 controls PAX6, we used the GRN to predict this regulation, and found that PAX6 might be indirectly down-regulated by OTX2 through DLX2 and POU2F2 (Extended Data Fig. 11i). Accordingly, DLX2 and POU2F2 regulomes are strongly affected by OTX2-LOF (Extended Data Fig. 11j). Altogether, these data bring together spatiotemporal GRN analysis with perturbations using genetic manipulation to highlight the utility of organoids and digital multimodal maps to gain holistic insight into human retinal neurogenesis.

## Discussion

Organoid models of human physiology and pathophysiology are becoming important multicellular systems for basic and translational research. However, the field has lacked integrative experimental and computational approaches for phenotyping organoid development across spatial and temporal scales. We show that 4i on tissues combined with single-cell genomics can be a flexible and broadly applicable approach to generate high-information content spatial and molecular descriptions of organoids and their primary counterparts. 4i on tissues is attractive as it utilizes off-the-shelf antibodies, and methods can be established for tissue processing, liquid handling, and confocal imaging that provide data spanning sub-cellular, cellular, and tissue scales. Here we generated a 52-plexed protein map from 41 samples, crossing length scales of ~150 nm to several mm in a total of more than 900 million multiplexed measurements. Our holistic analysis of organoid tissues provides data-driven approaches to explore global and local spatial heterogeneity. Thus, our multimodal map, together with previous assessments, suggest an optimistic view of the predictive capacity of retinal organoids. Indeed, we provide evidence that the well-known master regulator OTX2 is required to maintain retinal neuronal identities, consistent with results on non-human models (Ghinia Tegla et al., 2020; Nishida et al., 2003). The vast majority of chromatin access, gene and protein expression profiles, and cell differentiation profiles support the striking correspondence between organoid and primary retina counterparts. Human retinal organoids develop over many months in culture, and our data suggests that mosaic perturbation experiments can be performed at any point in development using inducible Cas9 iPSC lines to interrogate gene loss of function.

We also observe that organoids contain disorganized or malformed regions that are not often highlighted in the literature, and these deficiencies can hamper progress in translational research. By establishing a novel approach to assess intra- and inter-organoid heterogeneity based on tissue segmentation and clustering we can overcome barriers associated with organoid heterogeneity. We therefore expect that this approach will be useful for diverse organoid systems to explore tissue structure, understand tissue developmental dynamics, assess fidelity to primary counterparts, and quantify phenotypes in disease or other perturbation conditions.

## ACKNOWLEDGEMENTS

We thank the Camp, Treutlein, Pelkmans and IOB labs for helpful discussions. PW is supported by the European Molecular Biology Organization. GB is supported by the Engelhorn Foundation. JGC, BT, and LP are supported by CZF2019-002440 from the Chan Zuckerberg Initiative DAF, an advised fund of the Silicon Valley Community Foundation. JGC is supported by the European Research Council (Anthropoid-803441) and the Swiss National Science Foundation (Project Grant-310030_84795). BT is supported by the European Research Council (Organomics-758877, Braintime-874606), the Swiss National Science Foundation (Project Grant-310030_192604), and the National Center of Competence in Research Molecular Systems Engineering. LP is supported by the European Research Council (ERC-2019-AdG-885579), the Swiss National Science Foundation (SNSF grant 310030_192622), and the University of Zurich. JSF is supported by the Boehringer Ingelheim Fonds. We thank Josephine Jüttner and Özkan Keles for help with lentivirus preparation. Illumina sequencing was done by Ina Nissen, Elodie Vogel Burcklen and Christian Beisel of the Genomics Facility Basel; Genomics Core Facility (GeneCore) at EMBL Heidelberg, at the Single Cell Genomics Platform at IOB, Basel and at Novartis AG - NIBR - Chemical Biology & Therapeutics (CBT) Genomic Sciences. FACS sorting support was provided by Mariangela Di Tacchio, Aleksandra Gumienny and Thomas Horn of the single-cell facility at D-BSSE, ETH Zurich. The Botond Roska lab, Prisca Liberali lab and Lucas Pelkmans lab, shared many antibody aliquots and/or tissue samples. We thank Marietta Zinner and Urs Mayr for discussions of wet lab and analysis steps, Sandrine Bichet and Augustyn Bogucki of the histology facility of the Friedrich Miescher Institute for Biomedical Research (FMI) for help with histology work and use of their instruments. We thank Aaron Ponti, Erica Montani and Thomas Horn for assistance with microscopy, use of instruments and discussion of analysis steps.

## AUTHOR CONTRIBUTIONS

GB, AS, YH, PG, MR generated organoids used in this study. PW established, designed and performed the 4i experiments and developed and performed the preprocessing of the data, with support from GG, MH, LP, PN. GB, AS, DP, SP generated the single-cell transcriptome and accessible chromatin datasets. GB designed and performed the mosaic perturbation and Multiome experiments. CH, PW, PN, GG analyzed the 4i data. CH developed the Laminator and neighborhood analysis algorithms and performed related analysis with the support of ZH and PW. ZH, QY, BG integrated and analyzed the developmental time course multimodal single-cell genomic data. CH, ZH integrated 4i and single-cell genomic data to generate the digital retinal organoid. CH developed the web application EyeSee4is for visualization. JSF, ZH and GB analyzed the mosaic perturbation data. LP, MH, GG provided training and intellectual guidance for the 4i experiments and analyses. PW, GB, CH, ZH, BT, JGC designed the study, interpreted results and wrote the manuscript.

## DATA AVAILABILITY

The count tables generated in this study will be made available on Mendeley Data. Sequencing data will be deposited at the European Genome-phenome Archive (EGA). The data can be visualized in the EyeSee4is app (https://eyesee4is.ethz.ch/). Laminator and other codes used in the analyses will be deposited on GitHub (https://github.com/quadbiolab/).

## Methods

### Primary human tissue samples

Human retina tissue was obtained from a multi-organ donor as described in (Cowan et al., 2020). The sample included in this study is designated by the ID R-00646_06.

### Stem cell and organoid culture

We used four human induced pluripotent stem cell (iPSC) lines: B7 (01F49i-N-B7 (Cowan et al., 2020), IMR90 (iPS (IMR90)-4-DL-01, WiCell) and two lines expressing Cas9 under an inducible doxycycline cassette, B7 iCas9 (01F49i-N-B7-iCas9 with integration of the construct pAAVS1-ieCas9 described in (Ihry et al., 2018) in AAVS1 locus and 409B2-iCas9 (He et al., 2021)). For 4i experiments, B7 organoids at 6, 12, 18, 24, 39 weeks were used. Stem cell lines were cultured in mTeSR1 (Stem Cell Technologies, 05851) with mTeSR1 supplement (Stem Cell Technologies, 05852) or mTeSR Plus (Stem Cell Technologies, 100-0274) with mTeSR Plus (Stem Cell Technologies, 100-0275) and supplemented with penicillin/streptomycin (P/S, 1:200, Gibco, 15140122) on matrigel-coated plates (Corning, 354277). Cells were split 1-2 times per week after dissociation with EDTA in DPBS (final concentration 0.5mM) (Gibco, 12605010). Media was supplemented with Rho-associated protein kinase (ROCK) inhibitor Y-27632 (final concentration 5μM, STEMCELL, 72302) the first day after thawing. Cells were tested for mycoplasma infection regularly using PCR validation (Venor GeM Classic, Minerva Biolabs) and found to be negative. All cell lines showed normal karyotypes upon generation, the 409B2-iCas9 line acquired a common stem cell abnormality over time (He et al., 2021) and the B7-iCas9 acquired a duplication in the centromeric region of long arm of Chr20: q11.21-q11.22 (a known adaptation to cell culture growth conditions). To generate retinal organoids from the B7 and IMR90 line, 600 cells were seeded in each micro-well of an agarose chamber (MicroTissues® 3D Petri Dish® micro-mold Z764019) using the AMASS protocol (Cowan et al., 2020). 409B2-iCas9 organoids were generated with 1500 cells per micro-well and treated with BMP4 (55ng/ml, Biotechne 314-BP) on day 6 with half medium change on day 9, 12,15 (Kuwahara et al., 2015).

### 4i Sample preparation

Retinal organoids were embedded in 1% low melting agarose at ~37°C. The embedded organoids were then fixed in 4% PFA in PBS at 4°C overnight. Primary human retina samples were fixed in 4% PFA in PBS at 4°C overnight without agarose embedding. After fixation samples were transferred to 70% EtOH for at least 30 min before they were submitted to paraffin embedding which was performed according to standard procedure. A 96 well format cover slip (NEXTERION® Coverslips order ID 1535661) was coated with poly-l-lysine (SigmaP4832) for 1h at RT. Tissue sections were sliced on a microtome at 3μm thickness and transferred to the coated 96 well coverslip. The attached sections were then incubated at 37° C for at least 1h and then subjected to deparaffinization. Deparaffinization followed a standard procedure: 60° for 45’, 2 x 3’ Neoclear (Merck), 2 x 3 ‘ 100% ETOH, 1 x 3’ 96% ETOH, 1 x 70% ETOH, 1 x 5’ ddH2O. The samples were then fixed in 4% PFA in PBS for 15 min at RT and then subjected to antigen retrieval. Antigen retrieval followed a standard procedure. The sample plate was transferred to 800ml 1mM Citrate Buffer (pH 6.0, 0.05% Tween 20, sterile filtered) and heated to 100°C in a microwave (Milestone MicroMED T/T Mega) for 20 min. Subsequently the samples in the citrate buffer were left to cool to ~RT for ~2h with an open lid. The cover glass was then removed from the buffer and patted dry between the samples. Care was taken that the samples never dried out. The cover glass was then attached to the bottom of a 96 well self adhesive super structure (Merck GBL204969), placing each sample into a well.

### 4i imaging

The iterative staining and elution cycles were performed according to (Gut et al., 2018) but instead of an extra step for a 4’, 6-Diamidino-2-phenylindole (DAPI) staining Hoechst (Invitrogen 33342) was used and added to the secondary antibody staining solution at a dilution of 1:500. Pipetting was automated on a Hamilton Star robotic system. Secondary antibodies were applied at a dilution of 1:500. Primary antibodies and dilutions used can be found in Table S1. Imaging was performed using a Nikon Ti2 inverted microscope with a Yokogawa CSU-W1 SoRa spinning disk add-on. A 40x magnification was achieved using the Nikon PLAN APO 40x/1.25 SIL MRD73400 objective. The laser settings were kept constant throughout the experiment, 20% UV (405 nm), 20% green (488 nm), 20% red (561 nm) and 40% far red (640 nm) with 100 ms exposure each. Each two channels, 405 nm and 561 nm or 488 nm and 640 nm were acquired in parallel. The cameras used were Hamamatsu C11440 digital cameras. 20% of the sensors at each side was cropped to minimize shading effects and maximize stitching accuracy and 10% overlap was used for stitching. Images were acquired as z-stacks with a total of 6 μm thickness and 0.5 μm distance between image planes. Every image consisted of a 6×6 tiling array. The image cycling strategy constituted “staining cycles” (including elution and restaining) and “elution cycles” (including elution and mock staining steps without addition of antibodies). All cycles contained a Hoechst staining that was used for alignment of images across cycles. The elution cycles were performed every six cycles allowing assessment of background signal across the experiment (Table S1). The elution cycles were used for background subtraction (section “Background removal”). In every staining cycle three primary and secondary antibody combinations and a Hoechst stain were imaged. This was arranged so that always a rabbit AB was in green (488 nm) a mouse AB was in red (568 nm) and a third species antibody was in far red (647 nm) (Table S1). Several protein stains were excluded by visual examination (Table S1).

### Image preprocessing

A representative dark-field image was acquired by imaging 100 z-stacks with settings identical to image acquisitions but with lasers switched off. For each channel an average intensity projection per tile and an average of all tiles was generated resulting in the dark field images. Sample image tiles were maximum intensity z-projected (MIP) and for each channel the dark-field images were subtracted from every tile. No shading correction was performed. Stitching was performed after MIP using multichannel images with a Fiji plugin (Preibisch et al., 2009).

### Image registration

In every imaging cycle a Hoechst staining was included. These Hoechst images were used to align images across all cycles using SimpleElastix (Klein et al., 2010). The staining intensity of Hoechst significantly decreased over the rounds so that the signal between the early and late rounds was so different that not all cycles could be aligned to the first cycle. We therefore aligned images cycle00 - cycle06 to cycle01, cycle07 - cycle11 to cycle06, cycle12 - cycle16 to cycle11, cycle17 - cycle21 to cycle16 respectively. To restrict the alignment to the tissues of interest a simple initial tissue mask was generated that loosely defined the area of the tissue samples based on thresholding the Hoechst channel in the reference cycle. The 6×6 tiling array used for every sample did not always allow to include the entire tissue. To accommodate image boundaries that ran through the tissue the simple initial mask was updated to include only the area that was imaged in every cycle. Alignment was performed in two steps, a rigid transformation followed by an affine transformation. The parameters used for the elastix algorithm were optimized by visual examination of the results (Table S4). Images of every cycle and every sample were visually assessed for alignment accuracy. In cases in which alignment was suboptimal, different cycles were chosen as reference to improve the alignment.

### Image masking and denoising

After alignment we generated masks tightly surrounding the tissue samples per image to isolate the primary objects. To generate images that comprise reliable strong signal at the tissue periphery we merged the ATP1A1 (membrane) and the TUBA4A stains. To this end we scaled both images to the bottom 1st and upper 99th percentile, divided by 2 and summed the two images. The generation of masks based on these images utilizes the simple initial mask (as described in “Image registration”) multiple rounds of Otsu thresholding, dilation, expansion and closing of holes and adds a small rim of non tissue areas around the samples to ensure no tissue is excluded. Using these masks all images were then cropped to the bounding box containing the tissue. Images were then denoised using the denoise_nl_means algorithm from scikit image in fast mode (van der Walt et al., 2014). A sigma was estimated from the images. Further parameters were patch_size = 5, patch_distance = 6, cut-off distance (h) = 0.8 * estimated sigma. Several stainings had high spurious signals in areas corresponding to collagen labeled areas. These stainings were subsetted to exclude the staining overlapping the collagen stain. To this end a mask was generated from the collagen stains and the respective stainings masked using it. The pertaining Antibodies are marked in Table S1 and the images are flagged in the app EyeSee4is.

### Image background removal

We constructed images approximating the background signal per cycle by adding elution cycles proportionally for each staining cycle. We reasoned that the elution cycles represent the true background of staining cycles most closely in the cycles immediately following or preceding elutions. To account for the progressive shift of the background from the previous to the following elution in every set of five consecutive staining cycles we added the two elution cycles (before and after the stainings) proportionally. E.g. for staining cycle 11 elution cycle 10_0 and 15_0 were added at a 1:6 ratio respectively, for staining cycle 12 at a 2:6 ratio and so forth. To sum the elution cycle images both images were scaled to the upper and lower 1st percentiles, multiplied by the factor according to the desired proportion and summed. We then used a fast non negative least squares algorithm (https://github.com/jvendrow/fnnls) to gauge a factor with which to multiply the background model before subtracting it from the respective foreground image.

### Pixel matrix normalization

After registration every sample has a defined pixel grid that is consistent across all cycles. Therefore a matrix can be generated in which every row corresponds to a pixel and every column to antibody specific 4i imaging measurement in the same location. Imaging conditions were constant within every color channel throughout the experiment. To account for remaining signal divergences we normalized samples to one another while preserving divergent signal due to biological differences. We generated age specific expression matrices combining all images per age group. For every age group we calculated the mean and standard deviation per antibody channel. We then z-scored every image and multiplied it by the age and color channel specific standard deviation and added the age and color channel specific mean (reverse z-scoring) similar to (Gut et al., 2018). Finally every image channel was scaled to 0-1 where 0 is the bottom 1st percentile and 1 is the upper 99th percentile across all images.

### Multiplexed Tissue Units (MTUs)

Pixel clustering was performed similar to (Gut et al., 2018). In brief for both, single-well and global clustering image values were normalized and scaled as described in “Pixel matrix normalization”. For the global clustering the normalized global pixel matrix was randomly subsetted row wise by a factor of 1000 and submitted to FlowSOM (Van Gassen et al., 2015). FlowSOM was run using a 30×30 grid, Euclidean distance metric and 10 runs. The median intensities of the fSOM output clusters were further clustered using phenograph (Levine et al., 2015). The number of k-nearest neighbors specified for phenograph clustering was chosen by locating the inflection point on a graph plotting numbers of neighbors submitted to phenograph against number of clusters detected using KneeLocator of the kneed package (Satopaa et al., 2011). Subsequently all pixels were assigned to the phenograph clusters by closest euclidean distance in 4i intensity measurement space. For the single-well clustering the normalized pixel matrix was subsetted to include only pixels from an individual sample and then subsetted randomly row-wise by a factor of 1000. Each matrix was then submitted to FlowSOM, an inflection point was selected per sample as described for the global clustering and each sample was finally clustered using phenograph (Levine et al., 2015) with identical parameters as described above. For each sample all pixels were assigned to phenograph clusters using closest euclidean distance in 4i intensity measurement space. MTU images were generated by placing cluster assigned pixels into the masks outlining the organoid images and assigning colors to pixels according to the assigned cluster number.

### Nuclei segmentation and features

Nuclei were segmented using Cellpose (Stringer et al., 2021). We first thresholded the Hoechst signal of the reference cycle images by an Otsu threshold. We iteratively optimized Cellpose parameters for our samples. The parameters used were model_type: cyto, diameter: 35, flow_threshold: 0.8, cellprob_sizethreshold: 0, channels: [0, 0]. Nuclei features were retrieved using the segmented nuclei, the normalized pixel matrix (as described in “Pixel matrix normalization) and the regionprops_table function of the scikit-image toolbox (van der Walt et al., 2014) with standard scikit-image attributes or custom functions. Fluorescent intensity values measured per stain are bottom 5th and top 5th percentile, median, mean and pixel count (sum of all intensity values per nucleus). In addition spatial features, radial distance and nuclei density were measured per nucleus. For radial distance the refined mask was distance transformed and the mean distance value per nucleus was measured. For nuclei density an elliptical structuring element with 100*100 pixels (Bradski, 2000) was used as a kernel and the number of nuclei in a kernel around the centerpoint of each nucleus was used as a nuclear density measurement. For visualization nuclei features median fluorescence intensity values were scaled using the StandardScaler algorithm (Buitinck et al., 2013) and a UMAP embedding (Sainburg et al., 2021) was calculated from the first 8 PCs of the scaled features. Clustering was performed using the AgglomerativeClustering algorithm of the sklearn package (Buitinck et al., 2013) and the number of clusters was set to 8 for clustering of a single replicate. Nuclei density and radial distance values were assigned as raw values to nuclei in the UMAPs.

### Reconstruction of laminar organization dynamics

Contour outlines of all organoid samples and the adult primary sample were extracted using the skimage function find_contours() (van der Walt et al., 2014) after applying a gaussian filter (*σ*=50) and Otsu thresholding to the respective mask images. Note that the contour coordinates for the primary adult sample were filtered for only marking the outer circumference. Distance transformations were generated using the ndimage function distance_transform_edt() on respective mask images and applying a gaussian filter (*σ*=25). The center of the outer edges of the laminar windows (100 x 1000 pixels, 16.25 x 162.5 μm) were positioned on every 100th coordinate of the respective contour outlines and oriented along the inner-outer axis by iteratively rotating each window on their respective distance transformed image and maximizing the sum of captured distance transformation signal under each laminar window. Next, intensity profiles were extracted for all protein and MTU modalities by averaging across the inner-outer axis of each laminar window. Before analyzing the retained intensity profiles, laminar windows were filtered out that were positioned in straight parts of the contour outlines which are related to organoids that are not captured completely with the field of view. In addition, laminar windows that feature regions with no signal or damaged tissue were excluded by filtering windows for having more than 99% pixels with assigned MTUs. The laminar window intensity profiles of each protein and MTU modality were then reverse z-scored according to organoid section and age (as described in Methods Pixel matrix normalization) and scaled between 0 and 1. Before calculating distance matrixes for each feature, intensity profiles were smoothed by applying a 1D-mean filter (window size = 20) and downsampled by factor 2. Distances between laminar windows were then calculated by FFT-transforming the intensity profiles and calculating the euclidean distances of the first 10 frequency components for each feature utilizing the TSDist package in R (Mori et al., 2016). The resulting distance matrices were analyzed using the DistatisR method (Abdi et al., 2005). Laminar tissue heterogeneity was assessed by diffusion analysis (Haghverdi et al., 2015) of the *log*_10_(*x* + 1) transformed and across features aggregated distance matrix of laminar window intensity profiles. Results were then visualized by calculating UMAP embeddings for timepoint subsets of the first 10 diffusion components (DCs) and clustering laminar windows of each timepoint by performing Louvain clustering on the respective UMAP embeddings. In order to evaluate the clustering performance and properties of laminar window clusters, we reconstructed the MTU images for all oriented laminar windows and appended them ordered by time point-specific pseudotime. In addition, we mapped the detected Louvain clusters back onto Hoechst stain images of each organoid. We then manually selected laminar window clusters of each timepoint that did not feature RPE areas nor high within-cluster-heterogeneity. Note that due to the small number of laminar windows retained for the primary sample, we merged the respective Louvain clusters, thereby selecting all adult primary laminar windows for further analysis. A trajectory of laminar windows for the selected Louvain clusters (week 6: n=5, week 12: n=5, week 18: n=3, week 24: n=4, week 39: n=4, adult: n=1) were then reconstructed by again applying diffusion pseudotime analysis.

Similarity of laminar windows to the start and end point of the trajectory was assessed by calculating a maturation score. Therefore, the averaged compromised distances (gained from DistatisR analysis) to the upper and lower 5% quantile of pseudo temporal ordered laminar windows were calculated, subtracted from each other and scaled between −1 and 1. A constrained force-directed graph layout of laminar window Louvain clusters was generated by first filtering edges between clusters for being from the same or adjacent timepoints and then calculating edge weights by averaging the corresponding inter laminar window cluster compromised distances to construct a kNN graph (k=4) from the resulting adjacency matrix. Exemplary laminar window clusters for each timepoint were illustrated by randomly selecting 10 windows from each cluster and appending reconstructed MTU laminar windows along the pseudo temporal trajectory. For further visualization of the general abundance and patterns along the inner-outer axis of individual proteins and MTUs along the pseudo-temporal trajectory as heatmaps intensity profiles were processed as follows: Respective intensity profiles were scaled between their lower 1% and upper 5% quantiles and smoothened by position on the inner-outer axis along the pseudo-temporal axis by applying a 1D mean-filter (window size = 15). To visualize general abundance of proteins and MTUs along the pseudotime trajectory, intensity profiles were further averaged along the inner-outer axis and smoothed with another 1D mean-filter (window size = 25) along the pseudo-temporal axis in order to reduce noise in the generated heat map patterns.

### scRNA-seq, scATAC-seq and scMultiome for the developmental time course

Retinal organoids were generated from four different stem cell lines (B7, IMR90.4, 409B2-iCas9, B7-iCas9). 1-3 organoids of the same batch were pooled and dissociated at multiple timepoints of the course of retinal organoids development (Table S2). Organoids were washed three times with HBSS without Ca2+/Mg2+ (STEMCELL technologies, 37250). Single cell suspensions were obtained by adapting the protocol described in (Cowan et al., 2020) and using a papain-based dissociation kit (Miltenyi Biotec, 130-092-628). Briefly, 1ml of pre-warmed papain solution was added to the organoids and incubated for 10 minutes at 37°C. To facilitate dissociation, the mix was pipette-mixed every 5 minutes with a p1000. Enzyme mix A was then added and mixed by inversion and incubated for 10min at 37°C. At the end of the incubation, samples were pipette-mixed once more and successful tissue dissociation was confirmed via visual inspection. After incubation, cells were spun down 5min, 300g at 4°C. Cells were resuspended in 250μl of PBS+0.04% BSA and sequentially filtered through a 70μm filter (pluriSelect Mini 43-10070-50) and 40μm filter (pluriSelect Mini 43-10040-40). Cell counts were assessed with a Trypan Blue assay on the automated cell counter Countess (Thermo Fisher). For scATAC-seq, nuclei were isolated according to the protocol provided by 10x Genomics using the low input protocol and a lysis time of 3 min. Nuclei were loaded in a concentration that would result in a recovery of 10,000 nuclei. Single cell ATAC-seq libraries were generated with the Chromium Single Cell ATAC V1.1 Library & Gel Bead Kit. The libraries were sequenced on Illumina’s NovaSeq platform, NextSeq550 or HiSeq4000. For sc-RNA-seq, 8,000 cells were targeted and when the cell number was not sufficient, all cells were loaded. Single cell RNA-seq libraries were generated with the Chromium Single Cell 3’ V3.1 Library & Gel Bead Kit. Single cell encapsulation and library preparation were performed according to the manufacturer’s protocol. Libraries were pooled and sequenced on llumina’s NovaSeq platform, NextSeq550 or HiSeq4000. A summary of all single-cell experiments can be found in Table S2. Combined scRNA-seq and scATAC-seq (Multiome) were generated with the Chromium Single Cell Multiome ATAC + Gene Expression kit. Nuclei were isolated as described for scATAC-seq. The gene expression and accessibility libraries were FAB treated and sequenced on Illumina’s NovaSeq platform.

### Single-cell sequencing data preprocessing and multi-modal data integration

For the single-cell RNA-seq (scRNA-seq) data of each retinal organoid sample, Cell Ranger (10x Genomics, v4.0.0) with default parameters was used to align reads to the human reference (GRCh38, 10x Genomics, v.3.1.0) to generate the transcript count matrix for cells. Additional quality control (QC) was performed by excluding cells with detected transcript number fewer than 1500 or higher than 20000, as well as those with high mitochondrial transcript percentage (>20% for all the samples except >40% for GB2_scRNAseq). The exonic and intronic count matrices were generated via dropEst (Petukhov et al., 2018). For the single-cell ATAC-seq (scATAC-seq) data of each retinal organoid sample, Cell Ranger ATAC (10x Genomics, v1.2.0) with default parameters was used to align reads to the human reference (GRCh38, 10x Genomics, v1.1.0) to generate the peak fragment count matrix for cells. Additional quality control was performed by excluding cells with detected ATAC fragments fewer than 200 or more than 10000, cells with less than 20% fragments in the called peaks, cells with fragments at the blacklist genomic areas >2.5%, cells with nucleosome signal >3, and cells with transcriptional start site (TSS) <2.

To integrate the scRNA-seq and scATAC-seq data measuring the same cell suspension of a retinal organoid sample, a modified procedure of that as described before (Fleck et al., 2021) was used. In brief, we firstly used Seurat (v4.0.0) to normalize and log-transform the scRNA-seq data, identify highly variable genes (vst method, 3000 genes, followed by excluding mitochondrial and ribosomal genes), data scaling, principal component analysis (PCA, select top-20 principal components (PCs)), and Louvain clustering (resolution = 0.5). Average transcriptome profiles were calculated for each cluster. On the other hand, Signac (v1.4.0) was used to normalize the scATAC-seq data (default settings), perform partial singular value decomposition (SVD) on the normalized TFIDF of peaks with fragment detected in >0.5% of cells to obtain the latent semantic indexing (LSI) representation, and Louvain clustering (resolution = 0.5) based on the 2nd to 30th LSI components. Gene activity scores of annotated genes in each cell of the scATAC-seq data were calculated using Signac, and average gene activity scores were obtained for each scATAC-seq cluster. For each scRNA-seq cluster, the scATAC-seq cluster with the highest correlation across scRNA-seq highly variable genes which have non-zero gene activity scores in the scATAC-seq data was identified, and vice versa, resulting in a bipartite nearest neighbor network of clusters which contains multiple cluster components that are disconnected from each other. Next, canonical component analysis (CCA), implemented in Seurat, was performed on the scRNA-seq data and the scATAC-seq data represented by the gene activity scores based on the anchoring features selected by Seurat (same number as the highly variable genes in the scRNA-seq data). Lastly, a minimum cost maximum flow (MCMF) analysis was applied to cells from the two modalities in each cluster component. This constrained MCMF bipartite matching procedure resulted in bipartite edges each of which connects one cell in the scRNA-seq data with one cell in the scATAC-seq data. For each cell in the scRNA-seq data with at least one cell in the scATAC-seq data paired, a bi-modal metacell was formed with the RNA component being the scRNA-seq data, while the ATAC component being the union fragments of the paired scATAC-seq cell.

To evaluate the integration performance, an unweighted k-nearest neighbor (kNN, k=20) graph was firstly obtained for the scRNA-seq and scATAC-seq sample separately, based on Euclidean distances across top-20 PCs for the scRNA-seq data and Euclidean distances across the 2nd to 30th LSI components for the scATAC-seq data. We defined the distance between two cells as the shortest distance on the kNN graph, and then calculated the modal structure matching score (MSMS), which was defined as the Spearman correlation between the pairwise distances of cells in scRNA-seq with at least one cell in the scATAC-seq data paired, and the pairwise distances of the paired cells in the scATAC-seq as. A sample integration with MSMS<0.1 was considered as a failure, and were not considered when forming the bi-modal metacells.

For the scMultiome data, Cell Ranger ARC (10x Genomics, v2.0.0) was used to align reads of both the RNA library and ATAC library to the human reference (GRCh38, 10x Genomics, v2020-A-2.0.0) to generate both the transcript and peak fragment count matrices. Additional QC was applied with varied criteria for the two samples, both on the transcript count number, ATAC fragment number and mitochondrial transcript percentages. To generate a unified peak list to combine the chromatin accessibility profiles in different retinal organoid samples, we grouped all the scATAC-seq data into four groups based on organoid ages (0-10 weeks, 11-20 weeks, 21-30 weeks, >30 weeks), and used MACS2 to perform peak calling on each group separately and merge (using the Signac-implemented wrapper function CallPeaks in default parameters). Based on the new peak list, the fragment number per peak of cells in the scATAC-seq and scMultiome data were requantified.

To integrate all the single-cell data representing retinal organoids at different time points, we merged the data of the scRNA-seq (which contained both the ATAC-integrated metacells and cells without any paired cells in the scATAC-seq data) and scMultiome data mentioned above, together with a public scRNA-seq data of retinal organoids (Cowan et al., 2020). Focusing on the transcriptomic profiles, highly variable genes (vst method, 3000 genes) were identified for each of the three data sets, and those identified in at least two data sets were considered as the overall highly variable genes. Data scaling (cell cycle scores regressed out) and PCA (top-20 PCs) was performed, and the integration of different samples was done using Cluster Similarity Spectrum (CSS) (He et al., 2020) stratified on samples (cluster resolution=1). PCA (top-20 PCs) was then applied to the resulting CSS matrix to generate the PCA-reduced CSS representation. Louvain clustering (resolution=0.1) was applied, and one resulting cluster (cluster 6) was excluded for its expression of mesenchymal cell markers (e.g. DCN). Afterwards, the same procedure was applied to the remaining cells, resulting in the new PCA-reduced CSS representation of the data. Next, Louvain clustering (resolution=0.5) was performed. Cells in three of the clusters were further excluded from the following analysis, for their expression of brain cell markers (e.g. FOXG1, GFAP). The UMAP embedding of the remaining cells was generated given their PCA-reduced CSS representation.

In parallel, cells of all the scATAC-seq data, as well as those of the scMultiome data, were merged and integrated, considering their accessibility profiles across the unified peak list, using CSS stratified on samples, after data normalization with TFIDF and generating LSI representation with SVD (2nd to 30th LSI components) using peaks detected in >0.5% of cells. PCA was performed on the resulting CSS matrix to get the PCA-reduced CSS representation (top-20 PCs). Meanwhile, Harmony (Korsunsky et al., 2019) was also performed based on the 2nd to 30th LSI components with default parameters to generate a Harmony representation. Louvain clustering (resolution=0.5) was performed based on the PCA-reduced CSS representation, and the clusters with enriched cells without any paired cells in the scRNA-seq data (bonferroni-corrected one-sided Fisher’s exact test P<0.01) were excluded. Next, the cluster labels of the scRNA-seq cells (including the mesenchymal-like cells) were transferred to the scATAC-seq data for those cells with only one paired cell in the scRNA-seq data, or multiple paired cells but all share the same cluster label. For the rest of the cells, a network propagation procedure was developed to infer their corresponding cluster labels as follows. Firstly, three unweighted kNN (k=20) graphs were generated for the scATAC-seq cells, based on the LSI, PCA-reduced CSS and Harmony representation, respectively. The three graphs were averaged to form a weighted kNN graph. The adjacency matrix of the resulting kNN graph was obtained and row-normalized (i.e. sum of each row is 1). The normalized adjacency matrix, denoted as the propagation matrix P, was further modified, so that for any cell i which has a unique transferred cluster label from the scRNA-seq data, *P*_*i,i*_ = 1 and *P*_*i,j*_ = 0 if ij. This propagation matrix was then used to perform network propagation as *S*^*t*^ = *P* ∗ *S*^*t*−1^, where *S*^0^ is the initial transferred cluster identity matrix: 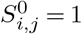 if cell i has a paired cell in the scRNA-seq data in cluster j, otherwise 0. The propagation was performed 100 times. The cluster with the highest propagated score was considered as the transferred cluster label of a given cell. The transferred-label-based cleanup was then performed by excluding cells with no paired cells in the scRNA-seq data, as well as cells with transferred cluster labels matching the mesenchymal or brain clusters. Afterwards, the similar procedure of dimension reduction and data integration across samples were applied. A UMAP embedding was generated based on the PCA-reduced CSS representation.

### Cell type annotation and developmental trajectory characterization

Based on the combinatorial expression of canonical cell type markers in the mentioned Louvain clusters (resolution=0.5), cells in the integrated cell atlas of retinal organoids in time course were firstly coarsely annotated as rods, cones, bipolar cells (BC), amacrine/horizontal cells (AC/HC), retinal ganglion cells (RGC), retinal progenitor cells (RPC), muller glia (MG), retinal pigment epithelial cells (RPE) and others. To further refine the annotation, we subsetted cells annotated as AC/HC, identified highly variable genes (default parameters) on the AC/HC subset, scaled the data, performed PCA (top-10 PCs), integrated data from different samples by MNN (Haghverdi et al., 2018) using the wrapper function in the R package SeuratWrappers with default parameters, and performed Louvain clustering on the MNN representation (resolution=0.5). Among those clusters, distinct amacrine cells (AC) and horizontal cells (HC) were identified. Similar procedure was applied to cells annotated as RGC. Among the Louvain clusters (resolution=0.3), cells were split into RGC and the earlier precursors. Afterwards, nine annotated cell types (RPC and eight terminal cell types rods, cones, BC, AC, HC, RGC, MG, RPE) were considered as well-defined cell types, while the rest of cells were considered as a pool of cells at varied intermediate cell states. The average expression profiles of the well-defined cell types were calculated, so as the normalized transcriptomic similarity of each cell to the nine cell types. This cell type annotation, as well as the normalized transcriptomic similarities, were transferred to the scATAC-seq atlas using the network propagation procedure as described above, based on a new weighted kNN graph based on the recalculated LSI, CSS and Harmony representation of the cleaned up cells in the scATAC-seq data.

Gene markers of each well-defined cell type were identified by comparing expression of cells of one cell type to cells of other well-defined cell types using the presto package in R combining multiple criteria (Benjamini-Hochberg (BH)-corrected two-sided Wilcoxon test P<0.01, AUC>0.7, average FC>1.2, detection rate difference>20%, being detected in fewer than 20% of the other cells, excluding mitochondrial and ribosomal genes), ordered by detection rates out of the cluster. To identify the peak markers of each cell type, a similar method was applied to the scATAC-seq atlas based on the transferred cell type labels. A putative marker peak was defined as one with BH-corrected two-sided Wilcoxon test P<0.01, AUC>0.51 and the ratio of detection rates in/out of the cluster >5, ordered by the detection rate differences.

To further investigate the cell state dynamics of retinal organoid development, scVelo (Bergen et al., 2020) was performed on the scRNA-seq data generated in this study, based on the stochastic RNA velocity model. The PCA-reduced CSS representation was used as the transcriptomic data representation. The RNA velocity transition matrix and velocity pseudotime were both obtained with default parameters. CellRank analysis (Lange et al., 2022) was then applied to the same cells using a hybrid kernel (50% velocity kernel and 50% velocity pseudotime kernel), with the eight terminal cell types considered as terminal states, to calculate the terminal states probabilities. The network propagation procedure as described above was used to transfer both the velocity pseudotime and the terminal state probabilities to other cells, based on the unweighted kNN (k=50) graph calculated from the PCA-reduced CSS representation.

Based on the information, we constructed a graph abstraction of the differentiation trajectories to different cell types from RPC as follows. Cells were grouped into three groups: AC, HC and others, and Louvain clustering (resolution=20) was applied to the PCA-reduced CSS representation of cells in each group to get high-resolution clusters, which were then pooled together, as the cell state representatives. The metadata of cells including both categorical (e.g. cell type annotation) and numeric (e.g. velocity pseudotime, CellRank terminal state probabilities) information were summarized to the clusters with either max-pooling or mean. PAGA, as implemented in scanpy, was used to compute connectivity between clusters given the PCA-reduced CSS representation (n_neighbors=20). Two clusters were connected if all the following three criteria were satisfied: PAGA connectivity > 0.2, one cluster being one kNN (k=20) of the other cluster in the summarized CellRank terminal state probability space, and the two clusters don’t belong to different terminal cell types. The connection was directional, from the cluster with lower pseudotime to the one with higher. A force-directed layout of the graph was generated for visualization.

To further refine the estimate of branching and trajectory towards different cell types based on the cluster-level abstraction, we chose five clusters with the lowest velocity pseudotime as roots and performed 100000 random walks starting from each of them until reaching a node with zero to-degree. Random walks ending at nodes representing non-terminal cell types were discarded. For random walks reaching each of the eight terminal cell types, the frequencies of passing by each node in the graph were counted and normalized by the number of random walks reaching that terminal cell type. For each node, the resulting normalized frequencies to different terminal cell types were further normalized by the sum to get the estimated likelihoods of the cell state being differentiating into different terminal cell types. Any likelihood <1% was set to zero and renormalized to get the final likelihood matrix.

### Reconstruction of gene regulatory network governing retinal organoid development

To reconstruct the gene regulatory network (GRN) controlling the retinal organoid cell type differentiation and maturation, we applied Pando (Fleck et al., 2021) to the integrated bimodal metacells together with cells in the scMultiome data. Pando incorporates evolutionary conservation, prior regulatory elements, data-driven open accessible regions (e.g. peaks called in the scATAC-seq data), TF motif database and binding site prediction to identify putative cis-regulatory elements and putative trans-regulators (i.e. TFs) of each gene, and fits a linear model of gene expression against interactions of the cis-regulatory element accessibility and trans-regulator expression, followed by statistical test to define significant TF-motif-target triplets. We applied Pando to the pseudocell-summarized data of the RNA-ATAC paired portion of the retinal organoid time course data to infer the GRN. The pseudocells were constructed by averaging transcriptomically similar cells from the same cluster of the same sample, following the procedure as described before (Kanton et al., 2019) and implemented in simspec package (He et al., 2020), with selection ratio of 0.1. This procedure was to decrease the noise due to the high sparseness of the data, especially the chromatin accessibility data. The extended TF binding motif database was constructed using similar procedure as described before (Fleck et al., 2021), integrating JASPAR (2020 release) (Fornes et al., 2019), CIS-BP database (Weirauch et al., 2014), and sequence-similarity-based motif transfer. Pando was run in default setting, except for the tf_cor threshold being 0.05. A TF-motif-target triplet was considered as significant if BH-corrected P<0.01.

To generate the layout of the resulting GRN highlighting TFs, the pairwise Pearson correlations of gene expression between different genes across the cell state representatives defined above were firstly calculated as the base TF-gene linkage score. Next, the lineage score between any TF-gene pair with no inferred direct regulatory relationship was set to 0. PCA (top-20 PCs) was then applied to convert the linkage score matrix to represent each TF, which was used as the input to generate the UMAP embedding of the TFs. For each TF in the GRN, its expression pseudotime was calculated as the average pseudotime of cell state representatives weighted by the expression level of the TF in different cell state representatives. The pseudotime dependency of the TF is calculated as R2 between its expression across cell state representatives and the predicted values of a smooth splines model of expression against pseudotime (degree = 3).

### Multimodal spatial integration

To integrate the spatially resolved time course of 4i nuclei with the multimodal scSeq dataset (Methods Section “Single-cell sequencing data preprocessing and multi-modal data integration”) we established an integration pipeline in Seurat (v4.0.0) in which we perform high resolution clustering in RNA space and transfer these meta cluster labels to the 4i data by CCA-anchoring. Protein stain intensity features of segmented 4i nuclei from the organoid samples of the developmental time course and the primary adult sample were *log*(*x* + 1) transformed and imported into individual Seurat objects. Note that we excluded several protein stains (Table S1) due to prior quality assessment as described above and protein stains that do not have a clearly associated gene in the RNA expression data (MAPK1/MAPK3, Serotonin, NPC and Hoechst). For each timepoint in the developmental time course (week 6, 12, 18, 24 and 39) the respective samples were integrated by CCA-anchoring using the Seurat functions FindIntegrationAnchors(dims=1:10) and IntegrateData(dims=1:10). Subsequently, PCA was performed on the integrated samples and UMAP embeddings were generated from the first 10 PCs for each timepoint. The adult sample was processed in the same manner but skipping the integration procedure.

We performed integration of the spatial resolved 4i nuclei datasets and the multimodal scSeq data for each timepoint of the 4i dataset separately by selecting respective matching and/or adjacent timepoints in the multimodal scSeq data. In order to maximize the proportion of ATAC-paired cells in the subsets we matched week 6 of the 4i data with week 6 of the multimodal scSeq data, week 12 with weeks 11, 12 and 13, week 18 with week 18, week 24 with week 24, week 39 with weeks 36, 38 and 40 and the adult sample with weeks 36, 38, 40 and 46, respectively. For each subset of the multimodal scSeq data Harmony integration (Korsunsky et al., 2019) was run accounting for sample grouping in order to evenly distribute paired ATAC cells among high resolution meta clusters which were subsequently computed with the Seurat function FindNeighbors(resolution=30, dim=15). In order to account for low total numbers of paired ATAC cells in matched subsets for week 24 and 39 we imputed ATAC signals for cells with no ATAC information within respective kNN graphs by applying network diffusion (n=20). RNA expression, ATAC peak regions and TF-motif score matrices were averaged by assigned meta clusters of each respective time point. In order to spatially resolve RNA expression, ATAC peaks, cell type specific ATAC peaks-sets and TF-motif scores, we transferred cell type and meta cluster labels from matched multimodal scSeq subsets to the corresponding 4i nuclei data subsets by CCA-anchoring. Note that for the visualization of the mapped “Intermediate” cell types in UMAP embeddings and onto organoid images, we calculated mixed colors based on the maximal correlation of intermediate cells with the defined terminal cell states for the multimodal scSeq data and further averaged these by meta clusters for visualizing 4i nuclei cell types. Furthermore, we added the position on the inner-outer axis of each nucleus to the metadata by calculating the euclidean distances to the respective organoid contour outlines. We also added the pseudotime rank of respective laminar windows as a binary count matrix which accounts for nuclei that might be positioned within overlapping neighboring laminar windows. For visualization of cell type and feature distributions along the inner-outer axis and pseudotime trajectory of laminar windows the integrated multimodal datasets were filtered for nuclei that occur at least in one laminar window with assigned pseudotime rank and then plotted as density contour estimates from pseudotime and inner-outer axis resolved cell type frequency or amplified feature count matrices. Count matrices for cell type abundances were generated by aggregating discretized nuclei positions along the inner-outer axis. In order to map continuous features like RNA expression or ATAC peak regions, count matrices of all nuclei were multiplied by their respective scaled (0-1) feature matrix, amplified by factor 10, rounded and aggregated.

### Spatial correlation and cell neighborhood analysis

To identify transcriptomic features and spatial cell type distributions that correlate with EPHB2 expression in week 39 organoids we focused our analysis on RNA transcripts that were variable expressed in the scRNA-seq data in week 38 and 40. Variable features were detected with the Seurat function FindVariableFeatures(). We then calculated two-dimensional kernel density estimations with an axis-aligned bivariate normal kernel for all respective per nuclei resolved spatial features across all organoid sections of week 39 using the kde2d() function from the R package MASS (Venables & Ripley, 2003) on a 200×200 grid. In addition, we calculated respective binary mask grids by calculating density estimations just based on nuclei positions and applying a threshold of 0.05. We then calculated the respective spatial correlations of all selected features with EPHB2 across all organoid samples after multiplying respective density estimations with their corresponding binary mask. Pearson’s correlations were then averaged by feature across all analyzed organoid sections. In addition, we calculated the spatial correlation of cell type distributions with EPHB2 in the same manner as described above. We then selected the top 115 positive and negative correlated features and ran GO term enrichment using the R package clusterProfiler (Wu et al., 2021) setting the background genes to the complete human genome. For visualization we plotted heatmaps of the top 4 positive and negative correlating features as well as EPHB2 in all analyzed organoid sections. Correlation of cell type distributions with spatial EPHB2 expression was visualized as a heatmap after applying hierarchical clustering (Ward.D2) of cell types across all analyzed sections. Results of the GO analysis were visualized as dotplots where we denote the detected gene ratios and −*log*_10_(*adjusted P values*). A complete list of input genes and resulting GO terms is shown in Table S4.

To analyze local, microenvironmental variation of RGC cells in the 4i data, we performed spatial resolved neighborhood analysis of protein and MTU signals within week 12 retinal organoids sections. We segmented RGC cell neighborhoods through a 40 pixel (6.5μm) radial extension from each nuclei centroid and averaged respective protein signals by radial position. Note that we masked respective nuclei in order to exclude protein intensity signals that stem from the nuclei themself and focus our analysis only on the respective radial neighborhoods. We then processed the retrieved radial intensity profiles according to Methods “Reconstruction of laminar organization dynamics”. In brief, we first applied z-scoring and scaled the intensity profiles between 0 and 1, smoothed the signals by applying a 1D-mean filter (window size = 20), downsampled by factor 2 and calculated respective distance matrices by applying FFT and averaging the euclidean distances of the first 10 frequency components. We then assessed the heterogeneity of RGC microenvironments by applying diffusion analysis, calculating UMAP embeddings from the first 10 diffusion components (DCs) and clustering radial neighborhoods by performing Louvain clustering on the UMAP embedding. Assigned Louvain clusters were then mapped onto respective week 12 organoid sections for visualization. We further averaged the MTU abundances within each Louvain cluster’s radial neighborhoods and applied hierarchical clustering (Ward.D2) and visualized the results in a heatmap. For further visualization of the detected radial spatial neighborhoods, we randomly selected 6 neighborhoods from all analyzed week 12 sections for each detected Louvain cluster and cropped respective image collages for Hoechst stain, all MTUs, a RGB overlay of POU4F2, HSPD1 and TUBB3 protein stains as well as a selection of MTUs 6, 8, 13,17 and 20.

As we observed RGC nuclei with apoptotic morphologies in several Louvain cluster neighborhoods, we further searched for apoptosis related processes in the scRNA-seq data of RGCs in weeks 11, 12 and 13. We therefore retrieved a list of human genes associated with the GO term apoptotic process (GO:0006915) from the AmiGO 2 database (http://amigo.geneontology.org/amigo/term/GO:0006915) which includes genes that are positively and negatively regulating apoptosis. We detected variable transcripts of RGCs by using the Seurat function FindVariableFeatures(). Next, we performed PCA for detected variable features that are also present in the retrieved list of apoptosis related genes and used the calculated PCA to run Harmony integration accounting for sample groupings. Based on the first 30 dimensions of the Harmony embedding we assigned clusters with the Seurat functions FindNeighbors() and FindClusters(resolution=1), and calculated a UMAP embedding. Gene markers of each detected RGC cell cluster were identified by comparing expression of cells of one cluster to cells of other cluster cell types using the presto package in R for all detected variable features. Cluster markers were selected by combining BH-corrected two-sided Wilcoxon test P<0.05 and AUC>0.5 criteria. We selected the top 50 markers from this ranking and used DAVID to perform GO term enrichment analysis and KEGG pathway analysis for cluster 6 which was defined by a set of marker genes strongly relating to apoptotic processes (see Table S5). We calculated a score for this detected apoptotic signature by using the Seurat Function AddModuleScore(ctrl=5) for the top 10 markers of cluster 6. For visualization we created feature plots of the UMAP embedding for the detected clusters as well as POU4F1, POU4F2, DDIT3, SCG2, ATF3 and apoptotic module scores of cluster 6.

### Vector and lentivirus preparation for perturbation experiment

The lentiviruses for perturbation experiment were produced according to (Datlinger et al., 2017; Fleck et al., 2021) with minor modifications. Briefly, a modified CROP-seq vector carrying GFP (Fleck et al., 2021) was used. Three gRNAs per targeted gene (NRL, OTX2, PAX6, VSX2, CRX) were designed using GPP Web Portal (https://portals.broadinstitute.org/gpp/public/) and synthesized by IDT following (Datlinger et al., 2017) recommendations. Moreover, a non-targeting ‘dummy’ guide (CGCTTC-CGCGGCCCGTTCAA) was added. Oligonucleotides were pooled in equal amounts and assembled in the vector backbone by Gibson’s isothermal assembly. The plasmid library was sequenced to validate the complexity of the pooled plasmid library and 10ng of plasmid library was used for generating a sequencing library with a single PCR reaction. Illumina i7 and i5 indices were added by PCR and the library was sequenced on Illumina’s MiSeq platform. Upon validation, lentiviruses were generated.

### In organoid CROP-seq perturbation experiment

Retinal organoids derived from 409B2-iCas9 line at 19 weeks of development were infected with the lentiviral pool described above. 20 organoids were individually infected in ultra-low attachment 96 well plates (Costar). Each organoid was infected with 1ul of LV (titer = 5.61E+08 TU/ml) in 100ul medium. After 4hrs, 150ul of medium were added for O/N incubation. The day after, organoids were moved to 10cm plates and treated with doxycycline 4ug/ml for 1 week to induce Cas9. After three weeks, 10 organoids were dissociated with the papain-based dissociation kit (Miltenyi Biotec, 130-092-628) described previously, and GFP+ cells were enriched by fluorescence activated cell sorting (FACS), (Experiment-1). The other 10 organoids were processed five weeks after infection (Experiment-2). We obtained a total of 12,000 and 7,000 GFP+ cells, respectively, and loaded them for scRNA-seq. We note that the first sample experienced a potential wetting error during GEM generation. Single cell RNA-seq libraries were generated with the Chromium Single Cell 3’ V3.1 Library & Gel Bead Kit. The expression libraries were FAB treated and sequenced on Illumina’s NovaSeq platform.

### gRNA detection from single cell cDNA

Guide RNA sequences from single cells were amplified from 30 ng of scRNA-seq cDNA as described in (Fleck et al., 2021). The first PCR amplifies a broad region targeting the outer part of the U6 promoter. The second, nested, PCR targets the inner portion of the U6 promoter adjacent to the guide sequence and adds a TruSeq Illumina i5 adapter. Finally, a third PCR adds Illumina sequencing i7 adaptors. PCRs were monitored by qPCR to avoid over-amplification. The samples were purified using SPRI beads (Beckman Coulter) and libraries were sequenced at 1:10 proportion of the transcriptome library on Illumina’s NovaSeq.

### CROP-seq data preprocessing and analysis

The reads of the CROP-seq experiment were aligned to the human reference (GRCh38, 10x Genomics, v3.1.0) with Cell Ranger (10x Genomics, v4.0.0) to generate the transcript count matrix for each sample. The amplicon sequencing reads detecting the gRNAs were aligned to the extended human reference (GRCh38-based, 10x Genomics, v3.1.0) using Cell Ranger (v4.0.0) with the artificial chromosomes representing the guide-GFP construct used in the CROP-seq experiment. Similar QC procedure as described before (Fleck et al., 2021) was used to extract informative gRNA transcripts detected in each cell based on a Gaussian mixture model of number of reads per UMI, resulting in a guide transcript count matrix for each sample, which was further binarized to obtain the final cell-to-gRNA and cell-to-target assignments.

The transcriptomic data of the CROP-seq experiment was merged and normalized using Seurat (v4.0.0). Highly variable genes were identified (vst method, 3000 genes, followed by excluding cell-cycle-related genes, mitochondrial and ribosomal genes), data scaling, PCA (top-20 PCs) were then performed, followed by data integration by CSS and further PCA reduction. To focus on retina relevant cells, the data was then projected to the CSS space (He et al., 2020) of the retinal organoid time course cell atlas described above. Based on the projected CSS representation, the kNN (k=50) of each cell (i) in the CROP-seq data in the time course cell atlas (denoted as 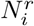), as well as the average of their distances in the CSS space (denoted as *d*_*i*_), were obtained. On the other hand, the average distance of each cell (j) in the time course atlas to its kNN (k=50) within the atlas 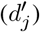 was also calculated. A normalized projected distance of each cell in the CROP-seq data to the time course atlas was thus defined as 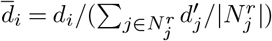. The bimodal distribution of 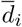 suggested a group of cells with large projection distance to the reference data which implied failure of the projection, i.e. cell types/states that did not exist in the reference data. A Gaussian mixture model was then fitted to identify those cells, which were excluded from the following analysis. Afterwards, a UMAP embedding was generated, and Louvain clustering with varied resolutions (0.2 and 0.6) was performed, both based on the PCA-reduced CSS representation of the remaining cells.

To estimate the perturbation probability of each cell being induced by the gRNA and Cas9 expression, we adapted the perturbation probability calculation as described (Fleck et al., 2021), using experiments and the Louvain cluster labels (resolution=0.6) as covariates. The resulting perturbation probabilities were considered as the proxy of perturbation status of a cell and used in the following differential expression (DE) analysis. The DE analysis was applied to co-expression gene modules, which were defined in the retinal organoid time course cell atlas as follows. A kNN (k=50) graph of genes detected in at least 1% of cells in the time course cell atlas was based on the Pearson correlation distance between gene expression across the abstracted cell state representatives. Louvain clustering (resolution=10) was applied to the gene kNN graph to identify the co-expression gene modules. The gene module activities were quantified for cells in the time course atlas, as well as cells in the CROP-seq data, separately, using the AddModuleScores function in Seurat with default parameters. An ANOVA-based DE method was then applied to the gene module activity scores of each co-expression gene module, testing whether the cell perturbation status of different targeting TFs significantly explained variance of the activity score of a gene module in the data set, with additional covariates such as experiments and cell clusters (resolution=0.6) included in the model. Co-expression gene modules with BH-corrected P<0.1 were considered as differentially expressed modules (DEMs) caused by the perturbation of the targeting TF. The size of the DE effect was represented by a signed −*log*_10_(*P*), where P is the ANOVA P-value, and the sign being the sign of the estimated coefficient. The resulting DEMs were clustered using hierarchical clustering (ward.D2 method), given the pairwise Pearson correlation distance across the time course cell state representatives. DAVID was used to do functional enrichment analysis of genes in each cluster of DEMs. The DE analysis was also applied to the two CROP-seq experiments separately with only the cluster label as the additional covariate, to estimate the robustness of the detected DE of the identified DEMs.

## Extended Data Tables

**Extended Data Tables**.

Table S1: Overview of 4i experiments

Table S2: Overview of single-cell genomic experiments

Table S3: Cell type marker genes and accessible chromatin

Table S4: EPHB2 spatially correlated genes GO term enrichment analysis

Table S5: Apoptotic RGCs GO term enrichment analysis

Table S6: SimpleElastix parameters

---

**Extended Data Fig. 1.**
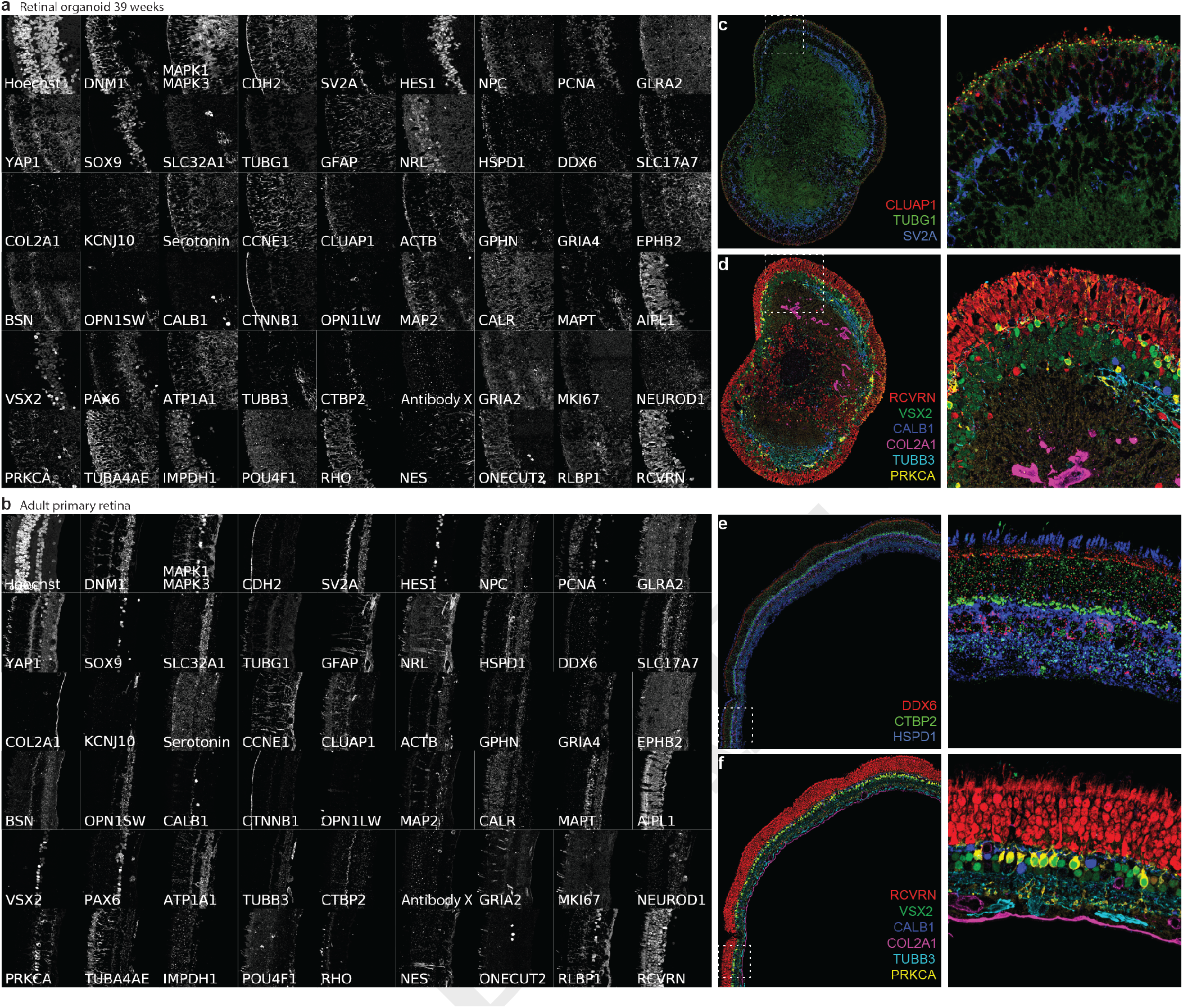
Representative images from primary adult human retina and a developed retinal organoid. a) Representative image segments from immunohistochemistry for each of the 53 antibody stainings and hoechst over 21 cycles from a human retinal organoid at 39 weeks (a) or primary primary retina (b). All images are from the same representative region of the organoid or the primary retina respectively. All images were scaled to the top and bottom 1st percentile except TUBG1 which was scaled to the top and bottom 0.1 percentile. c-f) Representative three (c, e) or six (d,f) way overlays of a whole 39 weeks organoid section (c, d) or a primary retina tissue (e, f). c) TUBG1 and CLUAP1 highlight centrosomes and ciliary bases or tips respectively. SV2A marks synaptic layers. d,f) RCVRN marks photoreceptors, VSX2 and PRKCA bipolar cell nuclei or rod bipolar cells respectively, CALB1 marks horizontal cells, COL2A1 marks the inner limiting membrane in the adult primary sample (f) or undefined collagen rich structures in the organoid section (d). TUBB3 marks neurons. e) DDX6 marks P-bodies, CTBP2 ribbon synapses and HSPD1 mitochondria.

**Extended Data Fig. 2.**
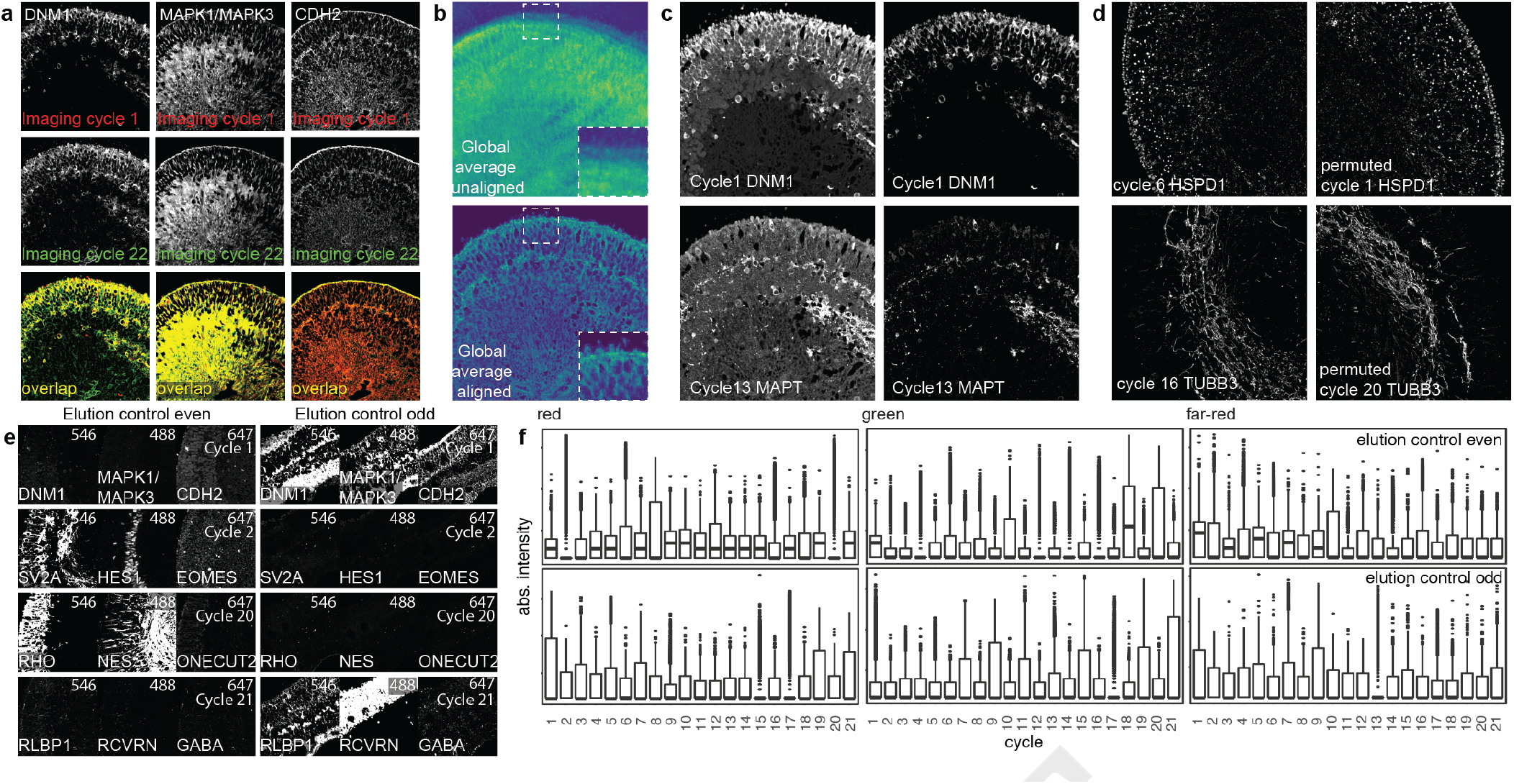
Quality control analysis of 4i data on tissues of developed retinal organoids. a) Images show the overlap (yellow) for three antibodies that were used for staining in cycle 1 (top row, red) and cycle 22 (bottom row, green) to assess restaining quality. b) Global average intensity signal of all stains (63 antibodies + hoechst) before alignment (top), and after alignment (bottom). c) Images of representative stains, cycle 1 DNM1 (channel 546) and cycle 13 MAPT (channel 647) before (left) and after (right) background subtraction. For representation, each image was scaled to the top and bottom 2nd percentile of the signal intensity. d) Effects of the antibody order was tested by perturbing the order in half the samples. Shown are two example AB stainings (HSPD1, top; TUBB3, bottom) in original (left; HSPD1 cycle 6; TUBB3 cycle 13) or permuted order (right; HSPD1 cycle1; TUBB3 cycle20). e) Examples of staining, elution, and re-staining efficiency. Two kinds of elution controls were included in the experiment. On the left control “even” was stained in even numbered cycles and “eluted only” in odd numbered cycles and on the right control “odd” was stained in odd numbered cycles and “eluted only” in even numbered cycles. In both cases the three immuno histochemical channels are represented (546 left, 488 middle, 647 right). All images were scaled to a constant absolute top and bottom value. Efficient elution was achieved until cycle 21. f) Signal changes over staining and elution cycles in elution control samples. The values were derived from fully processed (aligned, masked, denoised and background subtracted) images before exclusion of markers from the analysis. Masked images were randomly downsampled by a factor 1000 to select tissue pixels represented in the plots.

**Extended Data Fig. 3.**
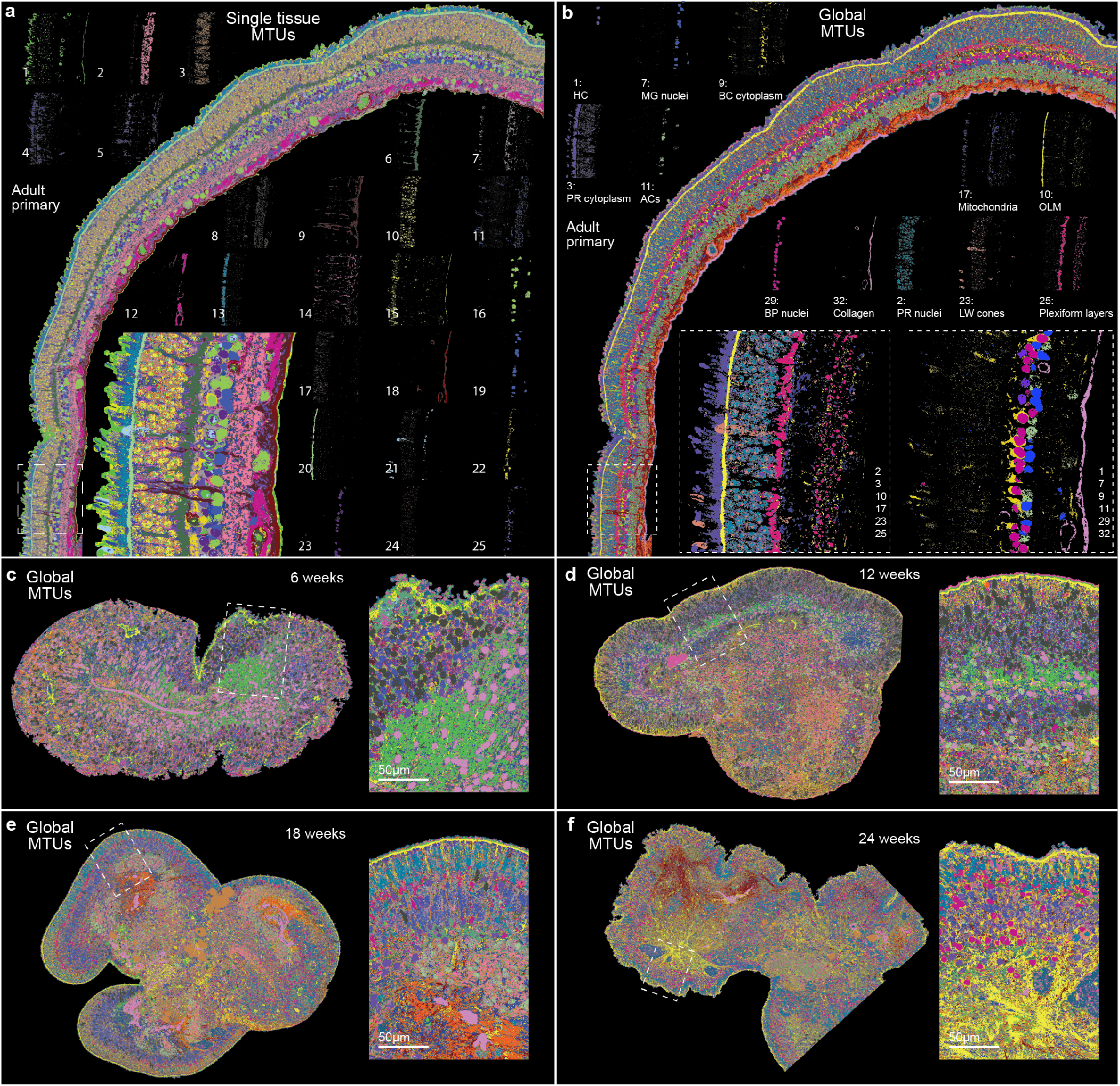
Assessment of single-sample MTUs in the adult retina and global MTUs across the organoid time course. a-b) Primary adult human retina section with pixels labeled by MTUs as generated by single sample clustering (a) or by global clustering including the entire organoid time course (b). Pixels assigned to each cluster are depicted individually for many of the clusters in each case. Highlighted clusters in b are the same as in Fig. 1e, f. Inset shows magnification of highlighted region (dotted line). c-f) Pixel assigned MTUs in organoid sections over the time course. MTUs were derived from global clustering and are comparable across samples including the 39 week samples shown in Extended Data Figure 3.

**Extended Data Fig. 4.**
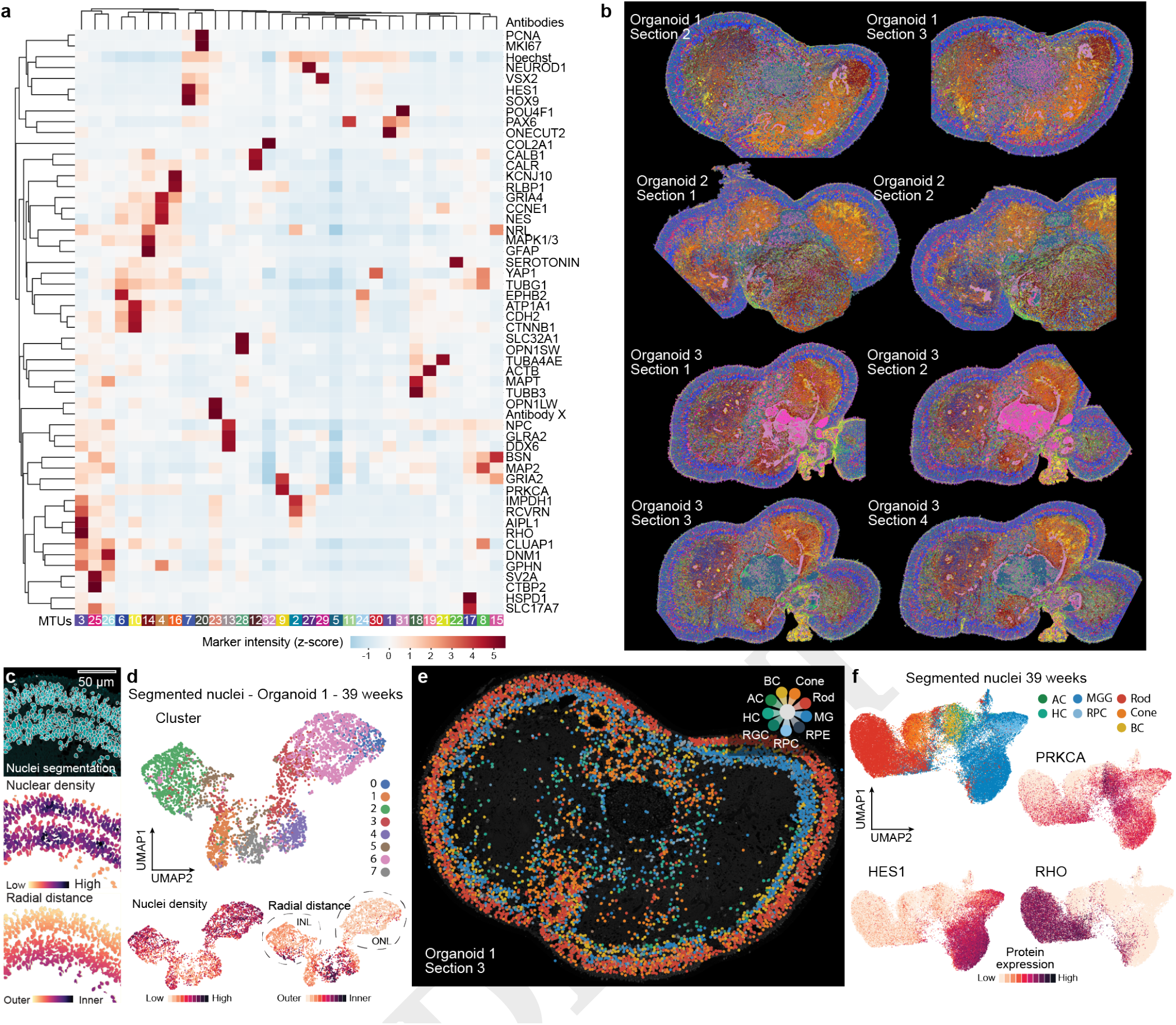
Extended analysis of 4i data on developed retinal organoids (39 weeks). a) Heatmap showing averaged protein detection intensity within each MTU. b) Representative MTU images from the three different 39 week organoids (6 out of 11 total sections analyzed from these organoids) show qualitatively congruent label assignment at a global scale. c) Nuclei segmentation allows measurement of features per nucleus. Features measured include spatial features such as nuclei density, radial distance and others or protein expression levels and MTU abundance measures. d) UMAP embedding of median immunohistochemical signal detected per nucleus of a single 39 week organoid section. Color coding based on agglomerative clustering resulted in 8 clusters (top panel) that broadly agree with cell types transferred from scRNA-seq data. Color coding of Nuclei density and Radial distance (bottom panels) highlight increased nuclei density in the INL and ONL and radial localization of the respective nuclei. e) Cell types transferred from scRNA-seq data mapped back into a representative 39 week organoid section. f) UMAP embedding based on protein features of all nuclei of 39 week organoid samples. Color coding based on cell types transferred from scRNA-seq data classifies nuclei into each of the major cell types. Signal intensity of HES1, RHO, and PRKCA highlight nuclei of the INL, ONL, and BC respectively.

**Extended Data Fig. 5.**
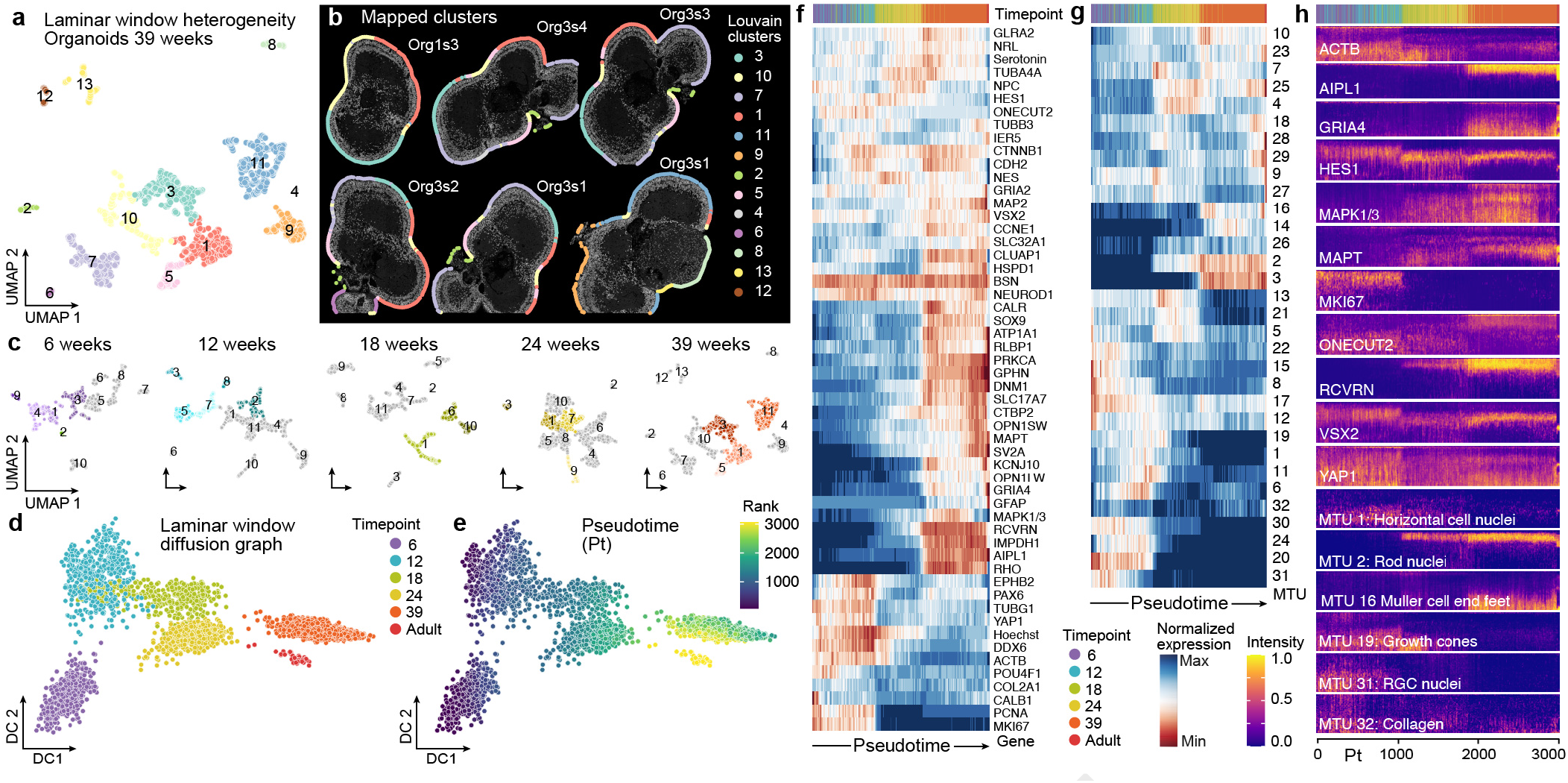
Extended analysis of laminar trajectory reconstruction. a) UMAP of laminar windows from all 39 week organoid 4i data colored and numbered by cluster. b) Image of each 39 week organoid section with window locations annotated as circles around the periphery and colored by cluster. c) UMAPs from the window heterogeneity analysis for each time point, with the clusters colored that were used for trajectory reconstruction. d-e) Windows projected onto diffusion component 1 and 2 from multidimensional diffusion analysis colored by time point (d) or by pseudotime rank (e). f-g) Heatmaps showing the average expression of protein stains (f) or MTU abundances (g) across windows ordered by pseudotime. Top sidebar colors each window by time point. h) Heatmaps showing protein expression or MTU intensities across laminar windows ordered by pseudotime, where each x position is a window with radial intensities averaged and then scaled across the inner-outer axis.

**Extended Data Fig. 6.**
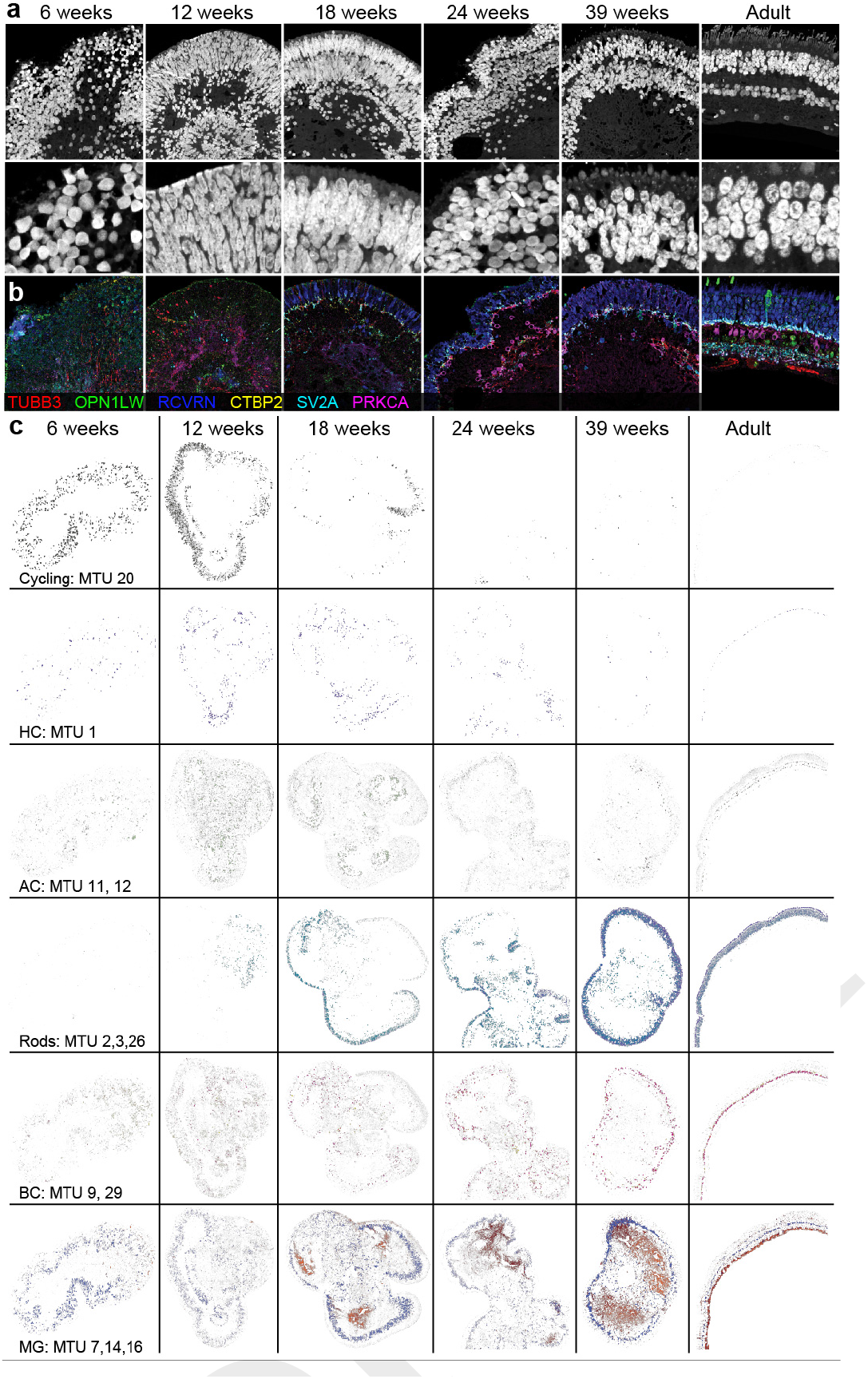
Extended Data Fig. 6. Analysis of features that change across the developmental time course. a) Hoechst stain in representative areas of organoids across the time course. Note the elongated nuclei at the periphery of the organoids at age 12 weeks and 18 weeks. b) Six color overlays in representative areas of organoids across the time course show emergence of synapses interfacing between neurons, BPs and PRs. TUBB3 (red) general neuron marker, OPN1LW (green) long wave cones, RCVRN (blue) photoreceptors, CTBP2 (yellow) ribbon synapses, SV2A (cyan) general synapse marker, PRKCa (magenta) rod bipolar cells. c) MTUs marking different cell states and types shown on organoid sections across the time course.

**Extended Data Fig. 7.**
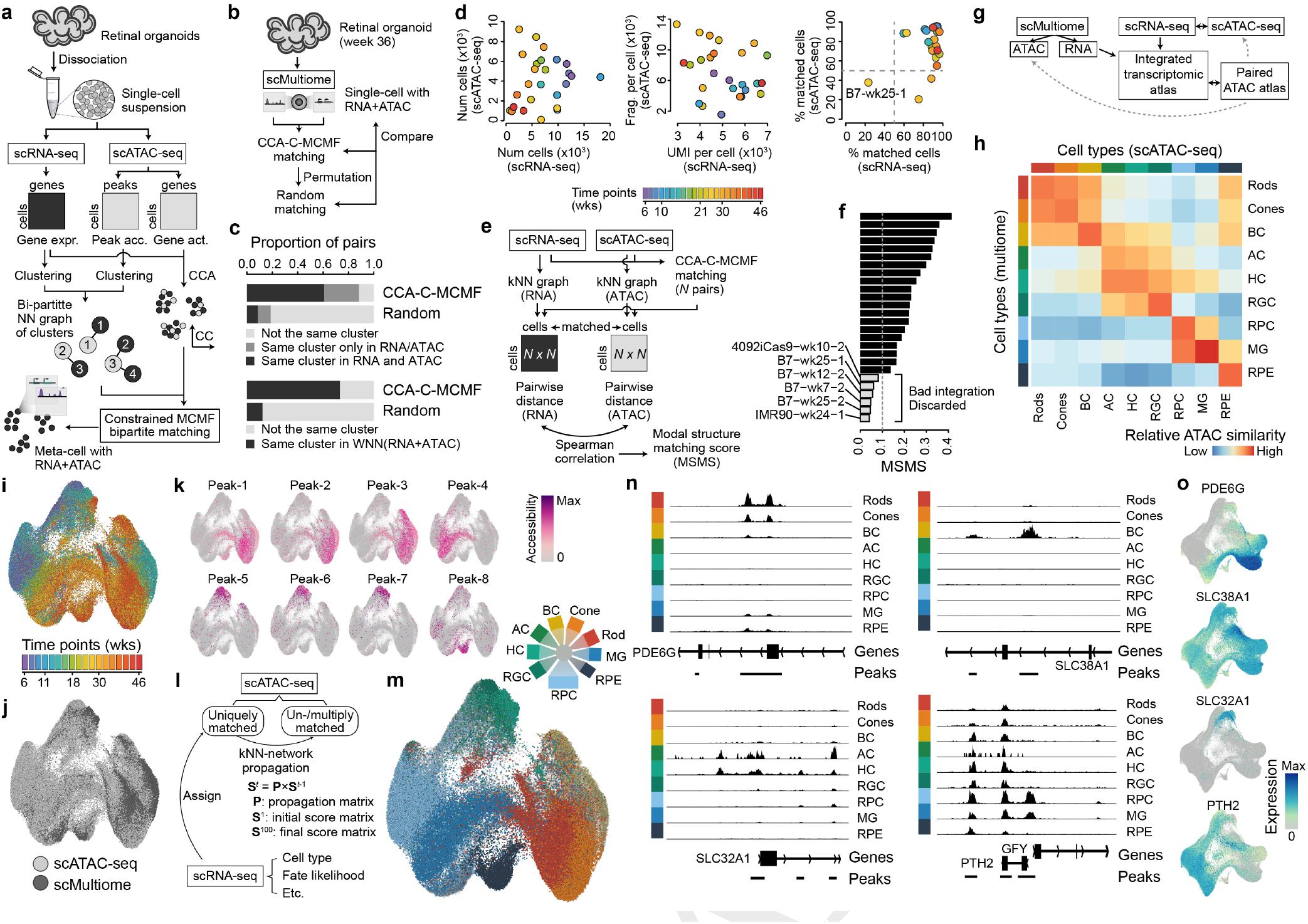
Supplemental analysis of multimodal integration and scATAC-seq data illuminates cell-type specific gene regulatory regions. a) Schematic of the experimental design and data integration strategy for the scRNAseq and scATACseq developmental time course. Organoids for each time point were dissociated and the cell suspension was used for scRNA-seq and scATAC-seq (see Table S2 for details). Modalities were integrated using minimum-cost maximum-flow (MCMF) bipartite matching in CCA space. The resulting metacells with RNA and ATAC components are integrated using CSS. b) Schematic outlining the integration strategy for multiome sequencing data where mRNA and chromatin access were measured in each single cell. c) Stacked barplots showing metrics assessing the metacell construction. d) Quality control assessments for the scATACseq datasets colored by time point. e) Schematic showing how multiome data was used to assess integration. f) Model structure matching score (MSMS) assessment of integration and rationale for sample discard. g-h) Schematic showing generation of the scSeq integrated multi-ome atlas, and heatmap showing the correspondence between cell types annotated based on scATACseq or multiome (RNA + ATAC, same cell) (h). i-k) UMAP embedding of scATACseq integrated with CSS and colored by time point (i), sample type (j), or by accessibility of cell type markers (k). l) Schematic showing the strategy for assigning labels to scATACseq cell types based on matching to cell types annotated from scRNAseq data. m) UMAP colored based on label matching from the scRNAseq data. n) Example of loci showing cell type-specific differential accessibility during retinal organoid development. Signal tracks show averaged chromatin accessibility in the different cell types. o) Transcriptome-based metacell UMAP colored based on genes shown in panel n.

**Extended Data Fig. 8.**
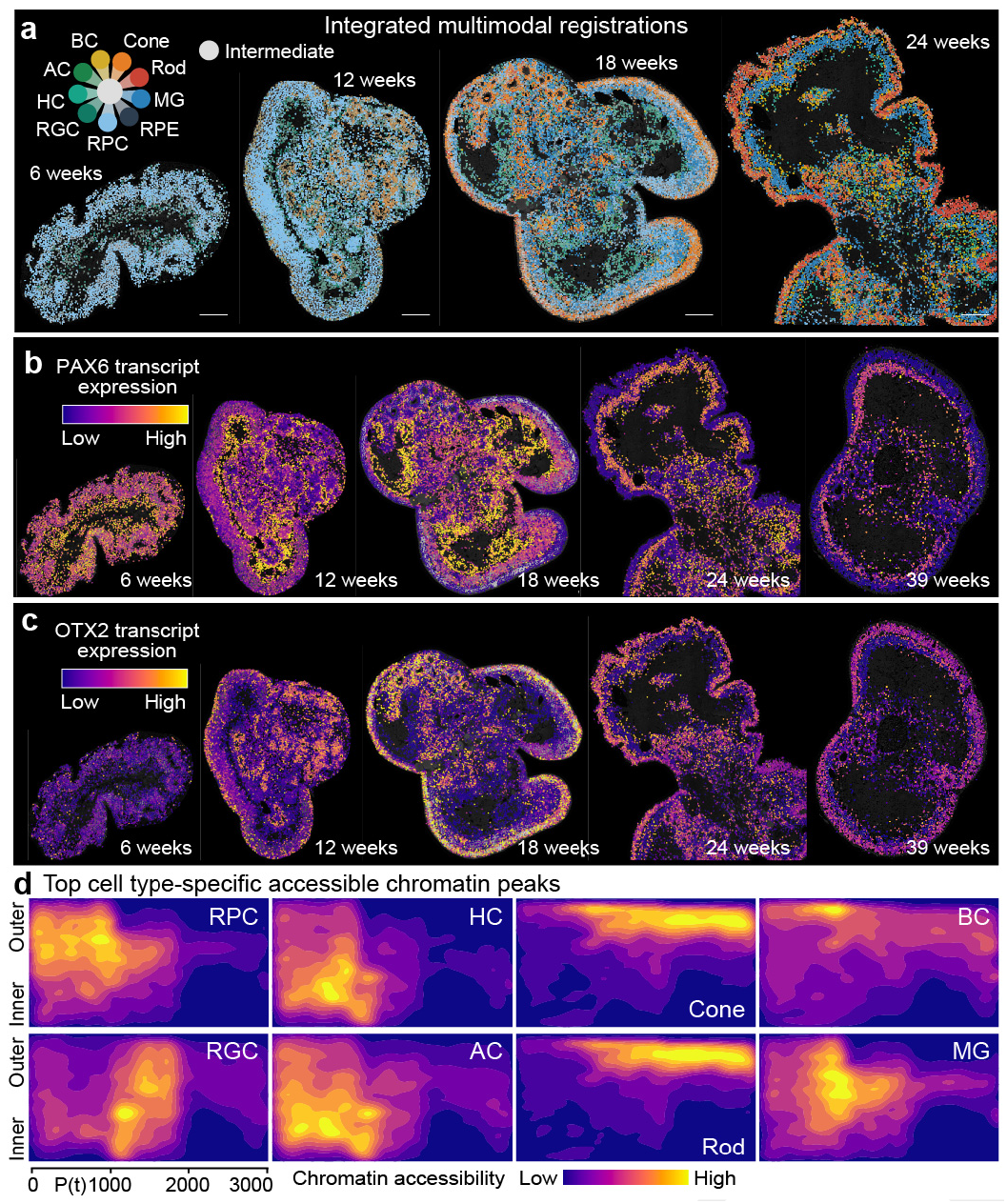
Integrated multimodal registrations with expression and chromatin access features across the retinal organoid time course. a) Overview of reference 6, 12, 18, and 24 week retinal organoid sections colored based on nuclei-type assignment from the label transfer. A corresponding 39 week organoid section is in Fig. 4b. b-c) Reference 6-39 weeks retinal organoid nuclei colored based on PAX6 (b) or OTX2 (c) RNA expression. d) Heatmap showing average chromatin accessibility densities of cell type-specific peaks for the major annotated retinal organoid cell types/states.

**Extended Data Fig. 9.**
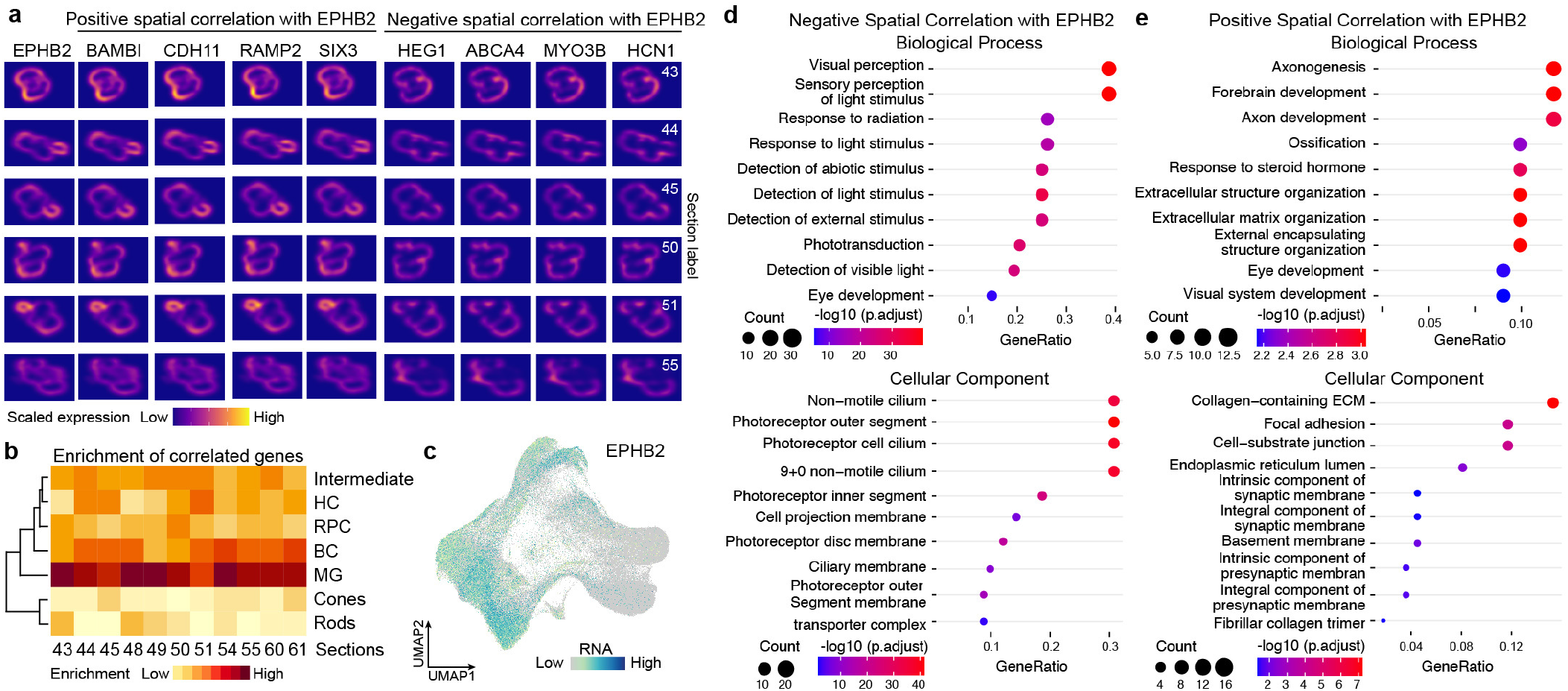
Integrated multimodal registrations with expression and chromatin access features across the retinal organoid time course. a) Overview of reference 6, 12, 18, and 24 week retinal organoid sections colored based on nuclei-type assignment from the label transfer. A corresponding 39 week organoid section is in Fig. 4b. b-c) Reference 6-39 weeks retinal organoid nuclei colored based on PAX6 (b) or OTX2 (c) RNA expression. d) Heatmap showing average chromatin accessibility densities of cell type-specific peaks for the major annotated retinal organoid cell types/states.

**Extended Data Fig. 10.**
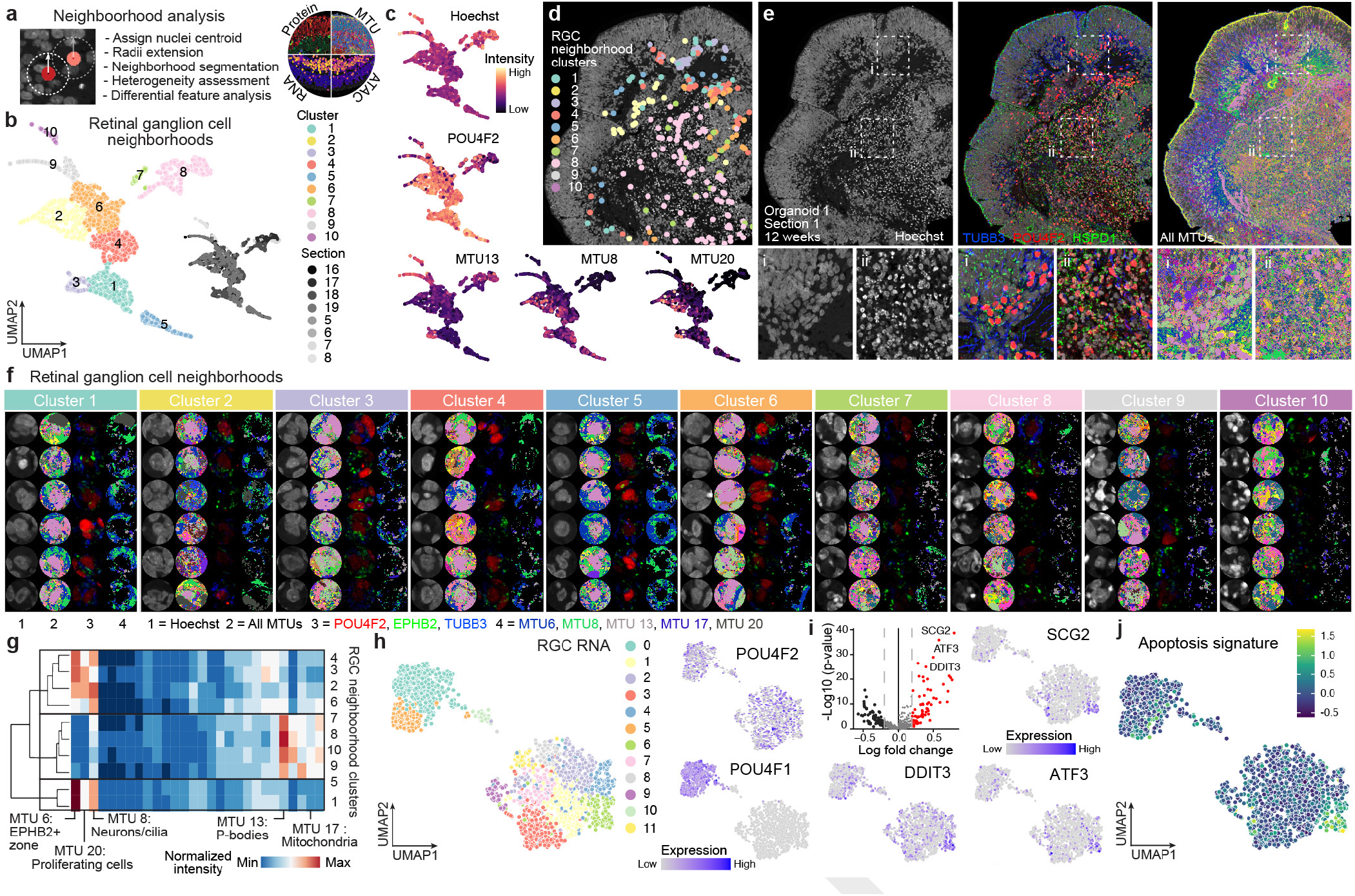
Identification of spatially-correlated transcriptome features associated with EPHB2 positive zones. a) Heatmaps from different sections of multiple 39 week organoids showing EPHB2 expression (left column) and genes that positively or negatively correlate with the EPHB2 spatial pattern. b) Heatmap showing spatial correlation of EPHB2 with each of the retinal organoid cell types across all week 39 sections. c) Feature plot showing expression of EPHB2 in the retinal organoid time course. d-e) Gene ontology Biological Process (top) or Cellular Component (bottom) enrichments for the top 115 genes that negatively (d) or positively (e) correlate with EPHB2 spatial domains.

**Extended Data Fig. 11.**
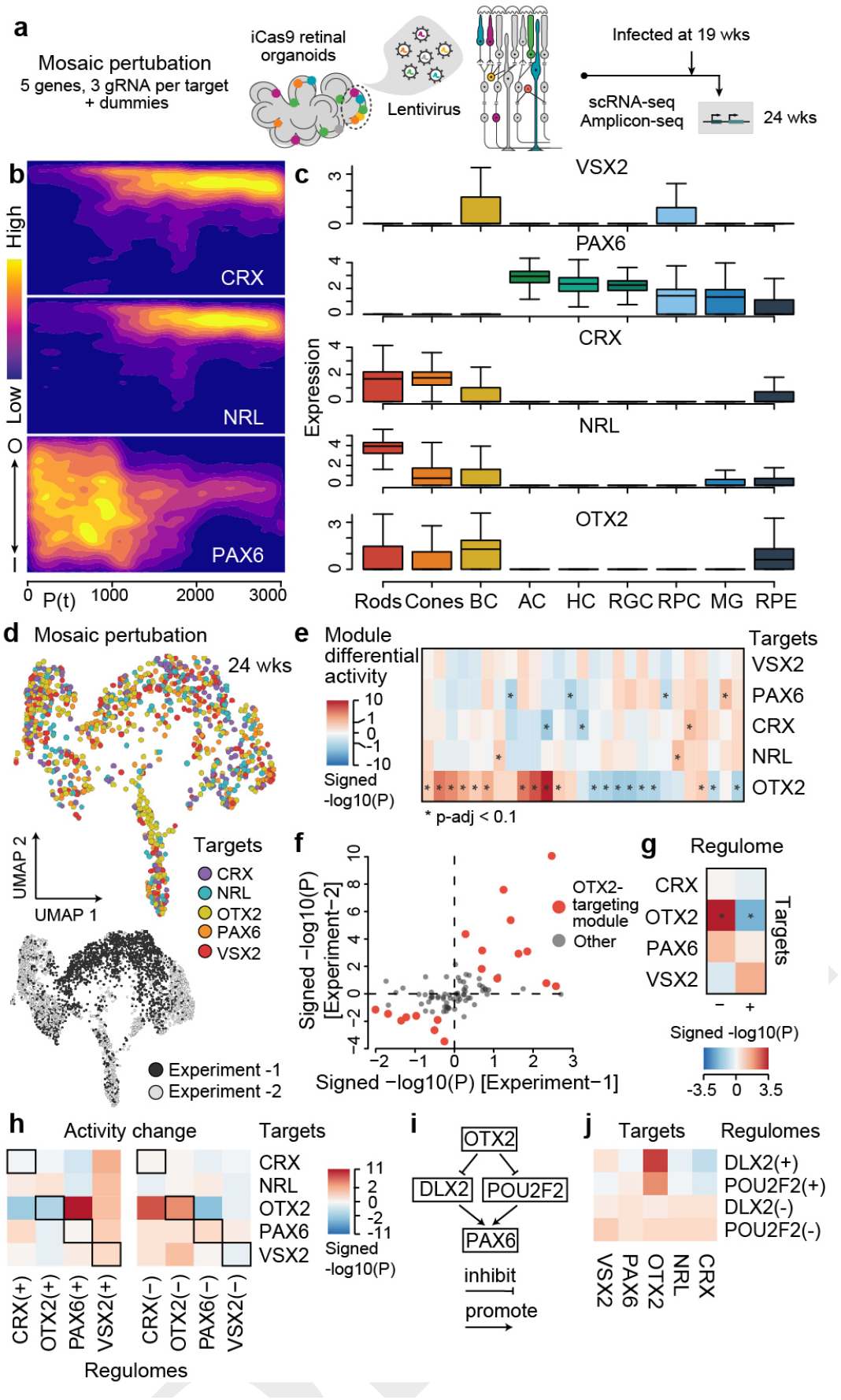
Analysis of retinal ganglion cell neighborhoods. a) Description of the neighborhood analysis algorithm. Cell neighborhoods were segmented through a 6.5μm radial extension from each nuclei centroid, and then searched for heterogeneity among neighborhoods based on annotated features (e.g. MTUs, protein, RNA, chromatin access). b) UMAP embedding of each RGC neighborhood colored and numbered by cluster or by organoid section (inset, greyscale). Embedding is based on variation in the MTU space. c) RGC neighborhood UMAP colored by various cluster features. d) A representative 12 week organoid with RGCs nuclei colored by cluster. e) Nuclei stain (left), immunohistochemistry (middle), and MTUs (right) of a representative 12 week organoid across the time course. POU4F2 expression (red) marks retinal ganglion cells (RGCs). Nuclei located towards the interior of the organoid are stained more intensely due to condensation and often fragmented, indications of apoptosis (nuclei inset). f) Six random exemplary RGC neighborhoods per cluster with nuclei, MTUs and protein stains highlighted. g) Heatmap shows hierarchical clustering of RGC clusters based on MTU intensities, with selected MTUs shown and provisionally labeled. h) RGC heterogeneity was analyzed in the transcriptome space. UMAP embedding shows RGC transcriptomes colored by cluster (left), or by RGC marker expression (right). i) Differential gene expression (DE) analysis between clusters and gene ontology enrichment revealed a signature (genes colored in red) in cluster 6 that is highly enriched for apoptosis biological process terms. The volcano plot shows the result of DE analysis between cluster 6 and other clusters, and feature plots show expression of top DE genes. j) RGC UMAP colored by the apoptotic signature (module score based on the expression of cluster 6 DE genes).

